# Phylogenomic evidence for the Origin of Obligately Anaerobic Anammox Bacteria around the Great Oxidation Event

**DOI:** 10.1101/2021.07.07.451387

**Authors:** Tianhua Liao, Sishuo Wang, Eva E. Stüeken, Haiwei Luo

**Author notes:** Corresponding author: Haiwei Luo, The Chinese University of Hong Kong Shatin, Hong Kong SAR. These authors contribute equally to this work (Tianhua Liao, Sishuo Wang).

## Abstract

The anaerobic ammonium oxidation (anammox) bacteria could transform ammonium and nitrite to dinitrogen gas, and this obligate anaerobic process accounts for up to half of the global nitrogen loss in surface environments. Yet its origin and evolution, which may give important insights into the biogeochemistry of early Earth, remains enigmatic. Here, we performed comprehensive phylogenomic analysis and showed a single origin of anammox bacteria within the phylum Planctomycetes. After accommodating the uncertainties and factors influencing time estimates, which includes implementing both a traditional cyanobacteria-based and a recently developed mitochondria-based approach, we estimated that anammox bacteria originated at early Proterozoic and most likely around the so-called Great Oxidation Event (GOE; 2.32 to 2.5 billion years ago [Ga]) which fundamentally changed global biogeochemical cycles. We further showed that during the origin of anammox bacteria, genes involved in oxidative stress, bioenergetics and anammox granules formation were recruited, which might have contributed to their survival on an increasingly oxic Earth. Our findings suggest the rising levels of atmospheric oxygen, which made nitrite increasingly available, was a potential driving force for the emergence of anammox bacteria. This is one of the first studies that link the GOE to the evolution of obligate anaerobic bacteria.

## Introduction

Anaerobic ammonium oxidation (anammox, NH ^+^+ NO_2_ ^-^→ N_2_ + 2H_2_O) [1], which usually occurs in anoxic marine, freshwater and wetland settings, accounts for up to 50% of the removal of fixed nitrogen (N) in nature. Along with denitrification, it is recognized as an important biological process that leads to N loss from the environment [2, 3]. In wastewater treatment, anammox is more cost-effective and environmentally friendlier than denitrification due to its lower oxygen requirement for aeration (nitrite [N(+III)], instead of nitrate [N(+V)] is sufficient for the anammox metabolism), carbon-free cultivation (some denitrifying bacteria are heterotrophic) and its negligible emissions of greenhouse gases like N_2_O [4, 5]. Consequently, anammox bacteria, the organisms that perform anammox, have been widely used in wastewater treatment plants [3, 6–8]. Despite the environmental and industrial importance of these organisms, their evolutionary history and antiquity are poorly known, which hinders accurate reconstructions of the biogeochemical nitrogen cycle over geologic time.

Previous studies investigated the roles of key genes driving anammox, including *hzsABC* (hydrazine synthase)[9], *hdh* (hydrazine dehydrogenase), *hao* (hydroxylamine oxidoreductase) [10] and *nxr* (nitrite oxidoreductase) [11]. In general, the vital enzymes encoded by these genes either are directly involved in anammox or participate in replenishing electrons to the cyclic electron flow [12, 13]. Further, all known anammox bacteria are found within the phylum Planctomycetes [14]. Six candidate genera of anammox bacteria, namely ‘*Candidatus* Brocadia’, ‘*Candidatus* Kuenenia’, ‘*Candidatus* Jettenia’, ‘*Candidatus* Scalindua’, ‘*Candidatus* Anammoximicrobium’ and ‘*Candidatus* Anammoxoglobus’, have been proposed based on 16S ribosomal RNA (rRNA) gene sequences [15], but none of them have been successfully isolated into pure cultures. The habitat of anammox bacteria requires the simultaneous presence of reduced (ammonia) and oxidized (nitrite) inorganic N compounds. Such habitats are often found at the aerobic-anaerobic interface in aquatic ecosystems, including the margins of oxygen minimum zones (OMZs) in the ocean and sediment-water interfaces, where ammonium originates from the anaerobic degradation of organic matter and nitrite can be produced by aerobic ammonia oxidation [16]. Ammonium, one of the two substrates of the anammox metabolism, was thermodynamically stable and likely present in moderate concentrations in the deep ocean throughout the Archean [17] and probably extending well into the Proterozoic [18]. The availability of nitrite, the second important substrate, is less certain. Geochemical data from sedimentary rocks suggest that the early Earth (before 3 Ga) was fully anoxic and deficient in aerobic ecosystems that were able to generate nitrite/nitrate [19–21]. The first transient and/or localized occurrences of aerobic nitrogen cycling appear in the Mid- to Neoarchean (2.7 Ga [22–25]), but evidence of widespread nitrate availability does not appear until around 2.4 billion years ago (Ga), the so-called Great Oxidation Event (GOE) [26–28]. The GOE marks the rise of free O_2_ in the atmosphere above 10^−5^ times modern levels, and was ultimately a result of the emergence of oxygenic cyanobacteria [29]. Hence it is conceivable that the appearance of anammox was linked to the appearance of aerobic ecosystems around the time of the GOE. However, it is also possible that some nitrite was provided much earlier, through lightning reactions in the Archean atmosphere [30]. Lightning, even in the absence of O_2_, can generate NO gas, which dissolves in water and converts into aqueous species, including nitrite [31]. Some anammox bacteria are even capable of using NO rather than nitrite directly as a substrate [12]. If these organisms capitalized on the lightning flux, then anammox might long pre-date the GOE.

Several tools exist for investigating the link between the evolution of metabolic pathways and geo-environmental transformations. One way to explore the evolutionary history of a specific metabolic pathway is based on organic biomarkers, which, however, are often affected by the poor preservation over geologic timescales [32]. For instance, ladderanes, a type of lipids that is unique to anammox bacteria [33], are rarely preserved to a level that can be used to date their evolutionary origin. Another approach is based on nitrogen isotope ratios of sedimentary records [17, 34, 35]. However, the isotopic fractionation factors for different metabolic pathways overlap widely (denitrification: −5 to −30‰ ; anammox: −16 or −24‰ [18, 36]), making it difficult to single out and elucidate the evolution of the different redox reactions within the N cycle by this method [37]. Alternatively, molecular dating, which estimates the age of the last common ancestor (LCA) of analyzed lineages by comparing their sequences based on the molecular clock theory [38], provides an alternative strategy to investigate this issue. Briefly, it can estimate the divergence timescale of organisms using genetic data while accounting for issues like uncertainties in the calibrations and different evolutionary rates among lineages, thereby bypassing the paucity and uncertainty of biomarkers and other biogeochemical records. On the one hand, because anammox is an anaerobic reaction, it is intuitive to assume that anammox should originate before the GOE, perhaps using nitrite or NO provided by lightning reactions in the atmosphere [39]. On the other hand, it is because of GOE that O_2_ on Earth rose permanently to a concentration that is biologically meaningful, and as a result, (micro-)environments were created in the surface ocean that were rich in nitrate/nitrite produced by aerobic organisms. These redox-stratified conditions may have stimulated the anammox metabolism by providing both nitrite and ammonium at the redoxcline. In this regard, it could be hypothesized that anammox did not arise until the GOE. Here, we ask the question when anammox originated. To answer this question and to obtain more insights into the evolution of anammox bacteria, we compiled an up-to-date genomic data set of Planctomycetes (see Supplementary Text section 1; Dataset S1.1), placed the evolution of anammox bacteria into the context of geological events, and investigated genomic changes associated with the origin of this ecologically important bacterial lineage.

## Results and discussion

### Monophyletic origins of anammox bacteria and anammox genes

The anammox bacteria form a monophyletic group within the phylum Planctomycetes in a comprehensive phylogenomic tree with 881 Planctomycetes genomes (Fig. S1). The anammox bacteria clade in this phylogenomic tree (Fig. S1) comprises four known genera including the early-branching *Ca.* Scalindua and *Ca.* Kuenenia, and the late-branching *Ca.* Jettenia and *Ca.* Brocadia. It also includes two understudied lineages which we named ‘basal lineage’ [40] and ‘*hzsCBA*-less lineage’ (Fig. S1), both of which have relatively low genome completeness compared to the other anammox bacteria and are represented by metagenome-assembled genomes (MAGs) sampled from groundwater (see Supplementary Text section 2.1). Note that the absence of *hzsCBA* in several genomes within the ‘*hzsCBA*-less lineage’ might be ascribed to the loss of these genes in evolution or their low genome completeness (Fig. S1). Because of their shallow (late-branching) phylogenetic position, this uncertainty should not affect the inference of the origin of anammox bacteria. Two described genera (*Ca.* Anammoximicrobium and *Ca.* Anammoxoglobus) do not have genomes available by the time of the present study (last accessed: April 2021), and are therefore not included in the phylogenomic analysis. Furthermore, using the last updated set of 2,077 Planctomycetes genomes (retrieved in April 2021) from NCBI, we obtained a consistent topology of anammox bacteria (Fig. S2).

To check whether important groups, particularly early-branching lineages, have been encompassed in our analyzed genome data sets, we built a 16S rRNA gene tree (Fig. S3) using the identified 16S rRNA genes from genomic sequences of anammox bacteria and the deposited 16S rRNA gene amplicons in SILVA from the class Brocadiae, to which anammox bacteria belong (see Supplementary Texts section 2.2). Since the anammox bacteria with genome sequences in this 16S rRNA gene tree (Fig. S3) showed a branching order congruent with the topology of the phylogenomic tree (Fig. S1), and since the 16S rRNA gene tree included 913 (clustered from 20,142) sequences sampled from a wide array of habitats (marine, sediments, man-made reactor, freshwater and terrestrial ecosystems), the 16S rRNA gene phylogeny, which could better capture the diversity of anammox bacteria by including uncultured samples, likely encompasses the early-branching lineages of anammox bacteria. Taken together, our genome set, which encompasses the genomes of the early-branching lineages, is appropriate to be used to study the evolutionary origin of anammox bacteria (Dataset S1.3). We further analyzed the evolution (Fig. S4) of the genes (*hzsCBA*) critical to the anammox reaction [10, 41]. Reconciliation (see Supplementary Texts section 2.3) of the gene and the genome-based species phylogenies pointed to a single origin of anammox metabolism at the LCA of anammox bacteria (Fig. S4).

Taken together, the above analyses suggest that our genome data sets have encompassed the earliest-branching lineages and deep phylogenetic diversity of anammox bacteria, and indicate its monophyletic origin, which agrees with previous studies [42–44]. This should provide a foundation for estimating the origin of anammox metabolism by estimating the origin time of anammox bacteria.

### The evolutionary origin of anammox bacteria coincided with the rising O_2_

We employed MCMCTree to estimate the divergence times of the anammox bacteria with i) soft bounds (time constraint allowing a small probability of violation) based on cyanobacteria fossils, and ii) the relaxed molecular clock approaches allowing substitution rates to vary among branches. We first employed a cyanobacteria fossil-based strategy, where cyanobacteria fossils were used as the only calibrations, a widely used strategy in dating deep-time bacterial evolution [45–47]. Using a best-practice dating scheme (Fig. 1; see Supplementary Text section 3), we dated the origin of anammox bacteria to 2,117 million years ago (Ma) (95% highest posterior density [HPD] interval, 2,002 - 2,226 Ma). Further, we repeated the analysis based on the expanded Planctomycetes genomes (Genome set 2; Dataset S1.1) and the constraint topology inferred with a profile mixture model (LG+C60+G; Genome set 1; Dataset S1.1). In general, both analyses based on different genome sets resolved consistent topologies (Fig. S5) as shown in Figure 1. Overall, they gave similar posterior times of the LCA of the anammox bacteria at 2,105 Ma (95% HPD: 1,961 - 2,235 Ma) or 2,005 Ma (95% HPD: 1,869 - 2,146 Ma).

**Figure 1.**
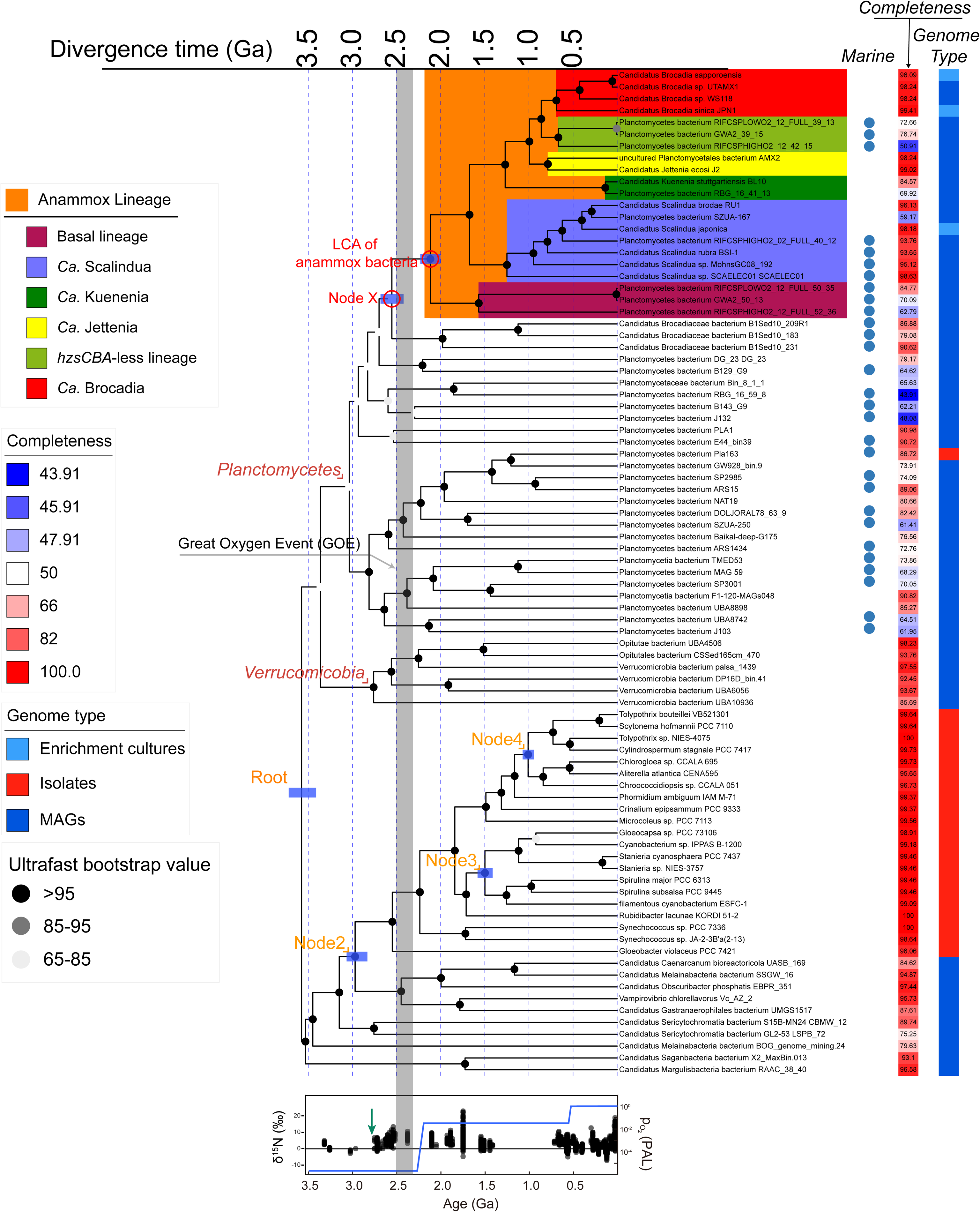
The evolutionary timeline of anammox bacteria using MCMCTree. The chronogram was estimated based on the calibration set C1 (see Supplementary Text section 3) and sequence alignments of the 25 orthologs conserved in bacteria (20040 sites). The blue bars on the four calibration nodes and the last common ancestor (LCA) of anammox bacteria represent the posterior 95% highest probability density (HPD) interval of the posterior time estimates. These alternative calibration sets were selected by choosing the three that accommodate different calibration constraints (see Supplementary Texts section 3.3). More alternative time estimates are provided in Figure S4. The vertical grey bar represents the period of the GOE from 2,500 to 2,320 Ma. The calibration constraints used within the phylum Cyanobacteria are marked with orange texts: the LCA of Planctomycetes and Cyanobacteria (Root), the total group of oxygenic Cyanobacteria (Node 2), the total group of *Nostocales* (Node 3), and the total group of *Pleurocapsales* (Node 4). Planctomycetes from marine or groundwater habitats were labelled by a blue circle. The genome completeness estimated by CheckM is visualized with a gradient color strip. The right next color strip indicates the genome type of genomic sequences used in our study including metagenome-assembled genomes (MAGs) and whole-genome sequencing (WGS) from either enriched culture sample or isolate. The diagram below the dated tree illustrates the change of atmospheric partial pressure of O_2_ (*P*O_2_) and nitrogen isotope fractionations. The blue line shows the proposed model according to Lyons TW, Reinhard CTPlanavsky NJ [29]. The green arrow suggests the earliest evidence for aerobic nitrogen cycling at around 2.7 Ga. PAL on right axis means *P*O_2_ relative to the present atmospheric level. The black dots denote the change of nitrogen δ^15^N isotope values according to Kipp MA, Stueken EE, Yun M, Bekker ABuick R [28].

Analyses with different combinations of calibrations and parameters broadly converge to similar time estimates (Fig. S6; Dataset S2.1). Specifically, assigning a more ancient time constraint to the total group of oxygenic cyanobacteria, which is based on the biogeochemical evidence of the presence of O_2_ at nearly 3.0 Ga [48] instead of GOE (Dataset S2.1), shifted the posterior dates of the LCA of anammox bacteria into the past by ∼300 million years (Myr) (e.g., C1 versus C10; Fig. S6). Varying the maximum bounds of the root by using 4.5 Ga, 3.8 Ga or 3.5 Ga had a minor impact (around 100 Myr) on the time estimates of anammox bacteria (e.g., C1 versus C3; Fig. S6). Though the auto-correlated rates (AR) model was less favored than the independent rates (IR) model based on the model comparison (see Supplementary Text section 3.3; Dataset S2.2), it is worth noting that the posterior ages estimated by the AR model were 1-12% younger than estimated by the IR model (Dataset S2.1).

While cyanobacteria fossils are the most widely used calibrations (and in many times the only calibrations) in bacterial molecular dating studies [47, 49, 50], there are two important issues: i) the available cyanobacterial fossils are very rare, ii) cyanobacteria fossils do not provide appropriate maximum calibrations [51]. Hence, we also used a recently developed mitochondria-based dating strategy [51] to estimate the origin time of anammox bacteria. This approach, based on mitochondria endosymbiosis, integrates mitochondrial lineages as a sister group to Alphaproteobacteria, thus taking the advantages of eukaryotic fossils, particularly several maximum calibrations, to improve date estimates. We compiled two gene sets (see Dataset S1.4), one according to the original 24 mitochondria-encoded genes used in [51] (mito24; 6295 sites), the other being six (mito6; 1238 sites) out of these 24 genes conserved across the bacterial tree of life (see Fig. 2A and Supplementary Text section 3.5). These analyses showed that the posterior time of the LCA of anammox bacteria shifted to the past by 90 to 500 Myr (Fig. 2), compared with only using the cyanobacteria-fossil-based approach (Fig. 1). The difference between calibrations C1 and C8 resulted from the removal of the maximum of the Node2 in C8, and accordingly, the posterior time estimated by C8 was greatly shifted to the past (Fig. 2B). When all maximum time constraints except for the one imposed on the root were removed, the posterior time was mainly affected by changes in the root maximum bound (C8 & C10; Fig. 2B). Despite the large phylogenetic distance between mitochondrial lineages and anammox bacteria, the mitochondria-based dating strategy may provide additional time constraints to the origin of anammox bacteria. For example, the ‘C8+Euk’ scheme effectively constrained the time estimates compared using a single maximum bound (C8; Fig. 2B). Furthermore, the posterior dates obtained by incorporating eukaryotic fossils (‘C8+euk’ to ‘C10+euk’) displayed less variation across calibration schemes (C8 to C10) based on cyanobacterial fossils (Fig. 2B). Nevertheless, the broadly consistent time estimates between analyses using cyanobacterial and mitochondrial calibrations highlight an origin of anammox bacteria roughly around or shortly after the GOE.

**Figure 2.**
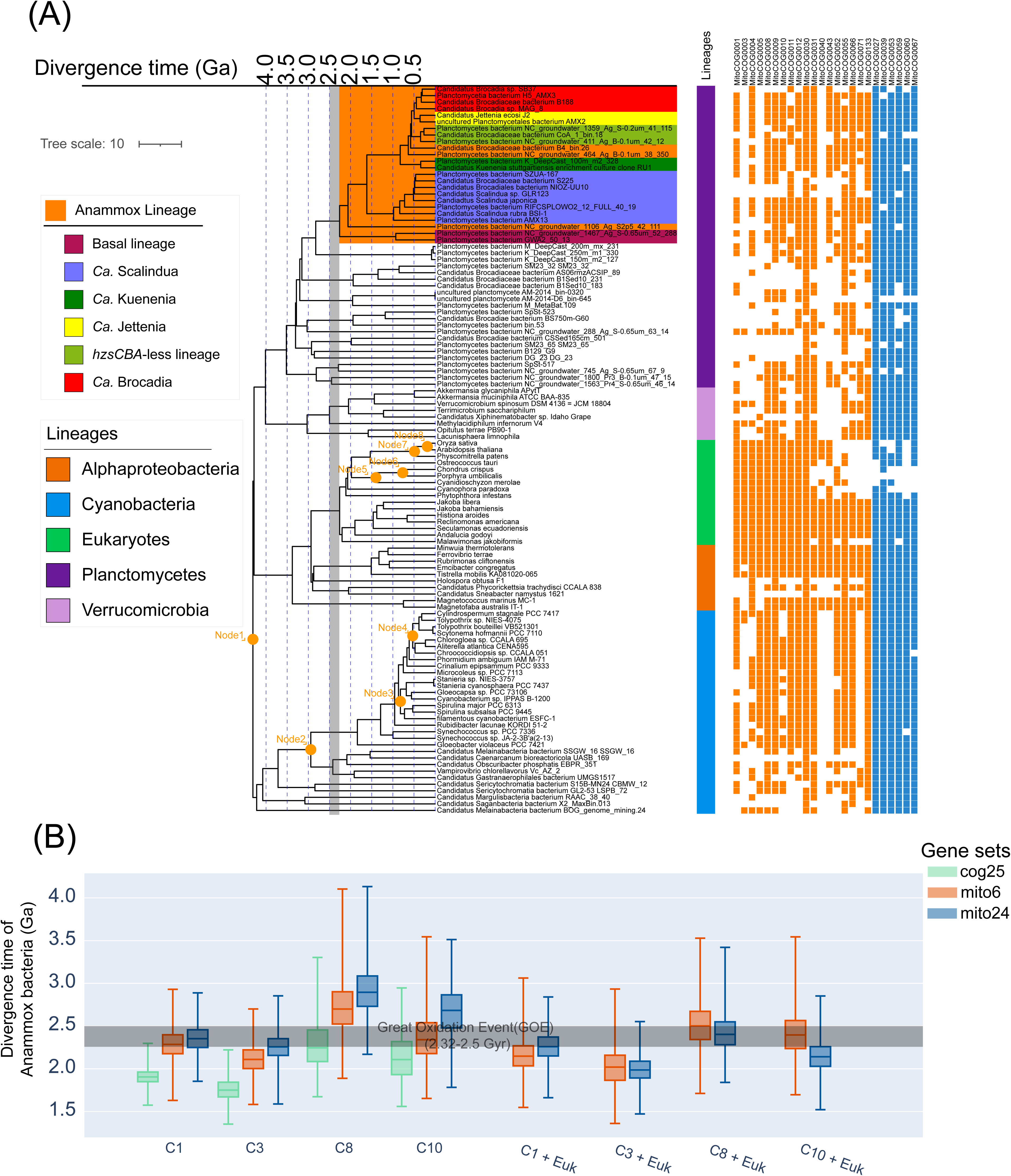
(A) The chronogram of the Anammox lineage estimated by mitochondria-based dating analysis. The time tree was estimated based on the calibration set “C1+Euk” and sequence alignments of the 24 mitochondria encoded proteins at the amino acid level (6295 sites). The vertical grey bar represents the period of the GOE from 2,500 to 2,320 Ma. The calibration constraints are marked with orange texts: the LCA of Planctomycetes and Cyanobacteria (Root), the total group of oxygenic Cyanobacteria (Node 2), the total group of Nostocales (Node 3), the total group of Pleurocapsales (Node 4), the total group of Bangiales (crown group of red algae) (Node 5), the total group of Florideophyceae (Node 6), the total group of mosses (crown group of Embryophyta) (Node 7) and the total group of eudicots (crown group of angiosperms) (Node 8). The color strip next to the label represent the major lineages of analyzed genomes. The filled and empty squares, respectively, represent the presence and absence of particular genes used in molecular dating analysis. The six genes selected to comprise the mito6 gene set are represented by blue squares. (B) The posterior times of anammox bacteria estimated using cyanobacteria-based and mitochondria-based dating analyses. The calibration sets starting with C represent the calibration sets with cyanobacterial calibrations, while those starting with Euk represent the calibration sets with eukaryotic calibrations. Calibration information: calibration scheme C1 node1: <4.5 Ga, node2: 3.0-2.32 Ga; C3 node1: <3.8 Ga, node2: 3.0-2.32 Ga; C8 node1: <4.5 Ga, node2: >3.0 Ga; C10 node1: <3.8 Ga, node2: >3.0 Ga. All of these calibration schemes share the same calibration for nodes 3 (>1.6 Ga) and 4 (>1.7 Ga). The horizontal grey bar represents GOE from 2.5 to 2.32 Ga. The detailed constraints of calibrations and time estimates are provided in Dataset S2.1.

Besides, there remains a possibility that there are unsampled or even extinct lineages of anammox bacteria that diverged earlier than all anammox bacteria analyzed in the present study. This scenario, if true, hints that the first anammox bacterium could have originated before the occurrence of the LCA of sequenced anammox bacteria but later than their split from the sister non-anammox lineage (Node X in Fig. 1; up to 2,600 Ma). Actually, a pre-GOE origin of anammox bacteria was also obtained by applying a larger minimum age (3.0 Ga) [52–54] to the total group of oxygenic cyanobacteria based on only cyanobacteria fossils (Fig. S5), or by using the mitochondria-based strategy (Fig. 2B). Accommodating the above uncertainties, our analyses suggest that the origin of anammox bacteria, and hence the origin of anammox, most likely falls into the 2.6-2.0 Ga interval, or more generally, around the early Proterozoic. Running MCMCTree analysis with no sequence data showed different distributions of time estimates for both the clade of anammox bacteria and the four calibration points (Fig. S7), suggesting that sequence data are informative for our molecular clock analysis [55]. Besides, we estimated similar origin times of anammox at around 2.1 Ga (the AR model) or 2.3 Ga (the IR model; see Fig. S9) with the complete taxon sampling shown in Fig. S2 (Genome set 2; see Dataset S1.1). For this analysis, we used a likelihood-based dating algorithm since the Bayesian molecular clock analysis is known to be computationally intensive and not suitable to take a data set with a broad taxon sampling [56]. Note that unlike Bayesian molecular dating software, the likelihood-based method we used only makes point estimations for the node ages (no standard deviation), thereby not allowing for fully appreciating the uncertainties in age estimates and any inference based on that. In general, the above result suggests that taxon sampling may not affect our dating analysis.

### The geochemical context of the origin of anammox bacteria

We speculate that the timing of the origin of the anammox metabolism is linked to the increasing availability of nitrite in surface environments, because ammonium was likely present in the deep anoxic ocean throughout the Precambrian [17, 18] and therefore presumably not a limiting substrate. The first abiotic source of nitrogen oxides on the early Earth, including NO and nitrite, would have been lightning reactions between N_2_ and CO_2_ in the Archean atmosphere [57, 58]. A prior study suggested that this process led to micromolar levels of nitrite in seawater [57], and the supply of NO may have been even higher, considering that it is the initial reaction product of lighting [57, 58]. This implies the possibility of an earlier origin of anammox bacteria. In fact, an intriguing hypothesis is that NO-dependent anammox [12] is ancestral to all anammox bacteria and arose before the appearance of significant nitrite levels with the GOE, perhaps driven by the lightning-source of NO. However, the original estimates of the lightning-derived flux of nitrogen oxides may have been based on perhaps unrealistically high amounts of CO_2_ [59]. Furthermore, to our knowledge, there has been no isotopic evidence in the early Archean rock record prior to 3 Ga for the presence of a significant nitrite or nitrate reservoir in the early ocean [19–21], and it is relatively conceivable that any nitrogen oxides that were supplied to the ocean by lightning were rapidly reduced to ammonium or N_2_ by ferrous iron, possibly even abiotically [60, 61]. Hence, although a background lightning flux of nitrogen oxides almost certainly existed, to our knowledge, there seems little evidence that it could supply a reliable metabolic substrate for early life. Furthermore, it is important to note that our phylogenetic results are not supportive of this hypothesis since the NO-utilizing anammox bacteria discovered to date are affiliated with phylogenetically shallow lineages from *Ca.* Kuenenia and *Ca.* Brocadia (Fig. 1). To our knowledge, there have been no studies reporting the ability of using NO as the sole electron acceptor in any other lineages of anammox bacteria. Hence, if NO-dependent anammox would have been present in the LCA of anammox bacteria, it was not clear why it was lost in all other anammox bacteria which resulted from multiple independent losses of the trait, a scenario apparently not favored from an evolutionary perspective (as shown by the arrows in Fig. S2). There are also anammox bacteria that are capable of using hydroxylamine as an oxidant; however, this molecule is a highly reactive compound and is usually considered as an intermediate in the nitrogen cycle, thereby rendering it a less important compound from the perspective of geological timescales. We are not aware of any non-biological sources of hydroxylamine. Taken together, our results are inconsistent with an Archean emergence of the anammox metabolism driven by lightning-derived nitrogen oxides. Instead, our results are most parsimoniously explained by the appearance of more significant nitrite/nitrate reservoirs around the time of the GOE.

The first transient appearances of nitrite/nitrate-dependent metabolisms are captured by the sedimentary nitrogen isotope record in late Mesoarchean soils at 3.2 Ga [23] and in Neoarchean shallow-marine settings at 2.7 Ga and 2.5 Ga [22, 24, 25]. These observations may reflect local and/or temporally restricted oxygen oases [62, 63]. Widespread nitrite/nitrate availability is inferred for the Paleoproterozoic (2.4-1.8 Ga), i.e., in the immediate aftermath of the GOE [26–28]. Importantly, O_2_ is necessary, although a trace amount is feasible, to extant nitrifying organisms for the oxidation of ammonium to nitrite and nitrate [64–66]. We noticed that a recent study [67] reports the ability of the model ammonia-oxidizing archaeal species (*Nitrosopumilus maritimus*) to continue ammonia oxidation after consuming all supplied O_2_ by nitric oxide disproportionation to generate O_2_, but the underlying genetic pathway is not known, making it difficult to extrapolate this mechanism to any other ammonia-oxidizing organisms. Hence the rise of oxygen almost certainly triggered the growth of the nitrite/nitrate reservoir, and therefore potentially provided one of the key substrates for anammox bacteria. We note that also NO would have become more abundant after the GOE, because it is an intermediate product in nitrification and denitrification [68]. This would put the possibility of NO-driven anammox back on the table. However, NO levels in seawater are highly variable, and so nitrite would likely have been a more reliable substrate. In any case, it would not violate our conclusion that the appearance of anammox in the Paleoproterozoic was linked to the GOE and directly or indirectly to the rise of marine nitrite. It has been shown that a low concentration of nitrite significantly decreases the rate of N removal by anammox bacteria in reactors [69, 70]. In modern OMZ in the Bay of Bengal, which may to some extent serve as analogues for the Paleoproterozoic redox-stratified ocean, the concentration of nitrite, instead of ammonium, was proposed as the rate-limiting factor to the anammox metabolism [71]. These imply that the rise of nitrite, which is driven by the rise of O_2_, could have facilitated the appearance of anammox bacteria in the early Proterozoic.

In modern environments, the majority of nitrite used in anammox is derived from nitrate reduction or aerobic ammonia oxidation [72], the latter performed by ammonia-oxidizing archaea (AOA) or bacteria (AOB). As noted above, nitrification could have appeared by at least 2.7 Ga, as indicated by nitrogen-isotope evidence for local accumulation of nitrate in surface ocean waters [18, 73, 74]. A recent study inferred that the LCA of AOA dates back to ∼2.3 Ga and first appeared on land, driven by increasing O_2_ concentrations in the atmosphere at that time, and that the expansion of AOA from land to the ocean did not occur until nearly 1.0 Ga [75]. If true, this hints at a dominant role of AOB in early ocean, before 2.3 Ga. Consistent with this idea, the vast majority of the early-branching lineages of anammox bacteria (*Ca.* Scalindua) and the sister Planctomycetes lineages of the anammox clade in the phylogenomic tree (Fig. 1) are found in marine or groundwater habitats. Furthermore, in the 16S rRNA gene tree which represents a greater diversity than the phylogenomic tree, marine lineages still account for the majority of the earliest-branching lineages of anammox bacteria (Fig. S3). Thus, it seems likely that the LCA of anammox bacteria originated in the marine realm where AOB thrived first. The above arguments, although speculative, tentatively suggest that the nitrite demand for anammox had been readily met by GOE by taking advantage of the significant amount of O_2_ newly available, which is broadly consistent with our molecular dating results suggesting an origin of anammox shortly before or after the GOE.

### Genomic changes potentially related to anammox metabolism and habitat adaptation upon the origin of anammox bacteria

We further explored the genomic changes characterizing the origin of anammox bacteria (see Supplementary Text section 4). This includes the gains of the aforementioned genes that directly participate in anammox, viz., *hzsABC*, *hdh*, and *hao*. The current view of the anammox metabolism posits that electrons consumed by anammox for carbon fixation are replenished by the oxidation of nitrite to nitrate by nitrite oxidoreductase (NXR) [13], which was inferred to be acquired upon the origin of anammox bacteria (Fig. 3). Moreover, multiple auxiliary genes for N assimilatory pathways including N regulatory protein (*glnB*), assimilatory nitrite reductase large subunit (*nirB*), transporters like *amt* (ammonium), *nirC* (nitrite) and NRT family proteins (nitrate/nitrite), were also gained at the origin of anammox bacteria (Fig. 3). However, genes for nitrogen-related dissimilatory pathways encoding nitrite reduction to nitric oxide (*nirK* or *nirS*) and to ammonium (*nrfAH*), respectively, were likely acquired after the origin of anammox bacteria (Fig. 3). Anammox occurs at the membrane of anammoxosome, an organelle predominantly composed of a special type of lipid called ladderane. The ladderane membranes show low proton permeability, which helps maintain the proton motive force during the anammox metabolism [33]. Although the biosynthetic pathway of ladderane is yet to be characterized, a previous study [76] predicted 34 candidate genes responsible for the synthesis of ladderane lipids, four of which were potentially acquired during the origin of anammox bacteria (see Supplementary Text section 4.2; Fig. 3), hinting at their potential roles in ladderane synthesis and providing clues to future experimental investigations.

**Figure 3.**
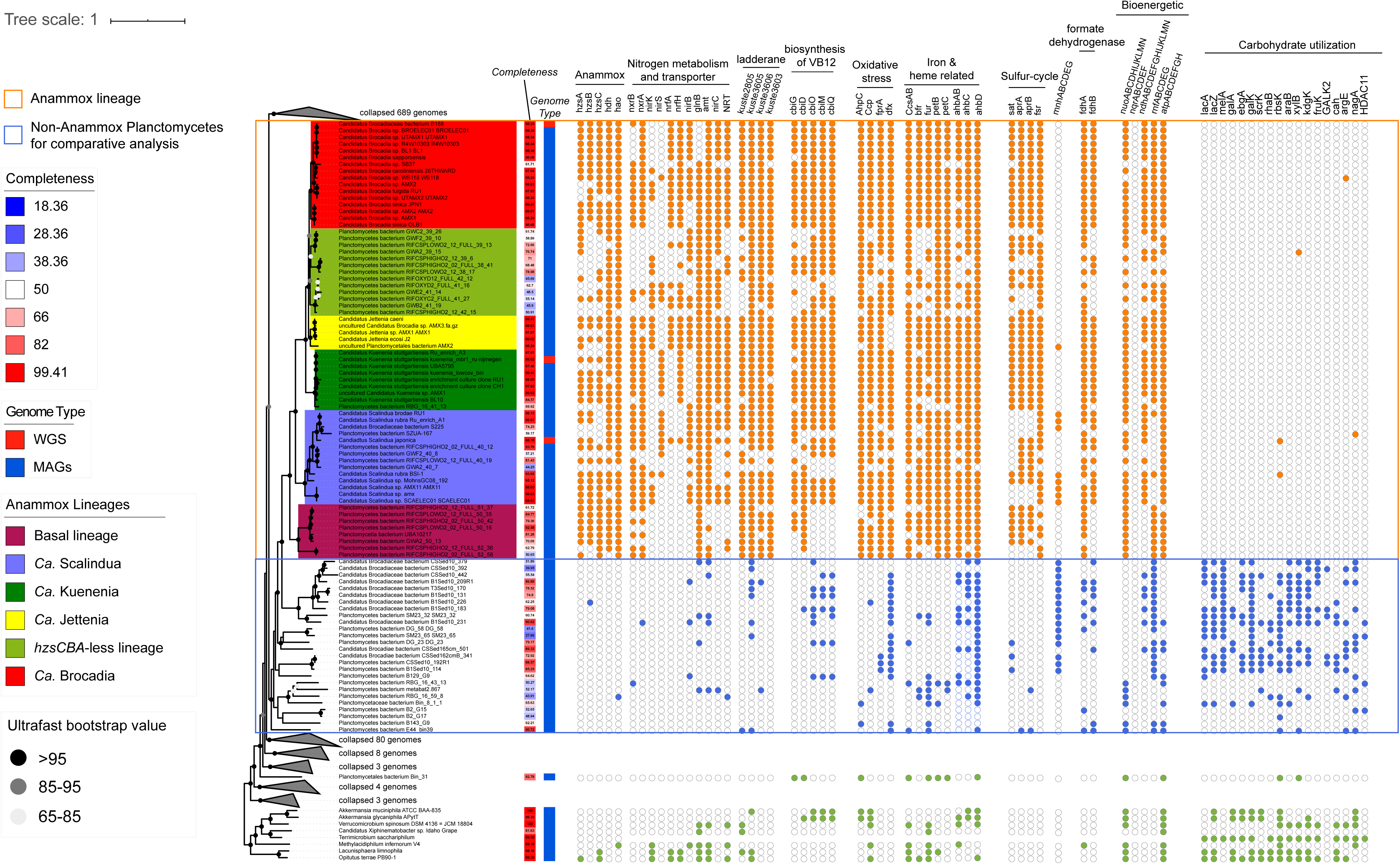
The phyletic pattern of ecologically relevant genes in the comparison between anammox bacteria and non-anammox bacteria. Solid circles at the nodes indicate the ultrafast bootstrap values in 1,000 bootstrapped replicates. Note that copy number difference is not indicated since most genes displayed here are present as a single copy in genome except a few exceptions like *hao* (see Dataset S1.5 for the table summarizing the copy number of each gene). The phylogenomic tree on the left was constructed with 887 genomic sequences described in Supplementary Texts section 2.1. Those Planctomycetes genomes not used for comparative genomics analyses are collapsed into grey triangles, and the numbers of collapsed genomes are labelled next to the triangles. The target group and reference group for comparative genomic analysis are within an orange or blue box, separately. For each genome, the genome completeness estimated by CheckM is visualized with a color strip and labelled besides leaf nodes. The right next color strip represents the type of genomic sequences used in our study including metagenome-assembled genomes (MAGs) and whole-genome sequencing (WGS) from either enriched culture sample or isolate. The filled and empty circles, respectively, represent the presence and absence of particular genes in corresponding genomes. For gene clusters, only genomes with at least half of the members of the gene cluster are indicated by a filled circle. The classifications of annotated genes are labelled above the gene names. *hzsCBA*, hydrazine synthase subunit C, B and A; *hdh*, hydrazine dehydrogenase; *hao*, hydroxylamine dehydrogenase; *nxrAB*, nitrite oxidoreductase subunit A and B; *nirK*, copper-containing and NO-forming nitrite reductase; *nirS*, cytochrome NO-forming nitrite reductase; *nrfAH*, ammonia-forming nitrite reductase subunit A and H; *glnB*, nitrogen regulatory protein P-II; *nirC*, nitrite transporter; NRT, nitrate/nitrite transporter; *amt*, ammonium transporter; kuste2805, 3603, 3605-3606, proposed genes relative to the synthetic pathways for ladderane at Rattray JE, Strous M, Op den Camp HJ, Schouten S, Jetten MSDamste JS [76]; *cbiG*, cobalt-precorrin 5A hydrolase; *cbiD*, cobalt-precorrin-5B(C1)-methyltransferase; *cbiOMQ*, cobalt/nickel transport system; *AhpC*, peroxiredoxin; *Ccp*, cytochrome *c* peroxidase; *dfx*, superoxide reductase; *fprA*, H_2_O-forming enzyme flavoprotein; *CcsAB*, cytochrome c maturation systems; *petB*, Cytochrome b subunit of the bc complex; *petC*, Rieske Fe-S protein; *ahbABCD*, heme biosynthesis; *sat*, sulfate adenylyltransferase; *aprAB*, adenylylsulfate reductase, subunit A and B; *fsr*, sulfite reductase (coenzyme F420); *higB-1*, toxin; *higA-1*, antitoxin; *mnhABCDEG*, multicomponent Na^+^/H^+^ antiporter; *fdhAB*, formate dehydrogenase subunit A and B; *nuo*(A-N), NADH-quinone oxidoreductase; *ndh*(A-N), NAD(P)H-quinone oxidoreductase; *nqrABCDEF*, Na^+^-transporting NADH:ubiquinone oxidoreductase; *rnfABCDEG*, Na^+^-translocating ferredoxin:NAD^+^ oxidoreductase; *atp*(A-H), F-type H^+^-transporting ATPase; *lacAZ*, beta-galactosidase; *melA*, galA, alpha-galactosidase; *ebgA*, beta-galactosidase; *galK*, galactokinase;, fructokinase; *fruK*, 1-phosphofructokinase; *rhaB*, rhamnulokinase; rbsK, ribokinase; *araB*, L-ribulokinase; *xylB*, xylulokinase; *kdgK*, 2-dehydro-3-deoxygluconokinase; GALK2, N-acetylgalactosamine kinase; *cah*, cephalosporin-C deacetylase; *argE*, acetylornithine deacetylase; *nagA*; N-acetylglucosamine-6-phosphate deacetylase; *HDAC11*, histone deacetylase 11.

Anammox bacteria occur in anoxic habitats and exhibit low oxygen tolerance [15]. Specifically, the O_2_-sensitive intermediates generated by anammox, e.g., hydrazine, a powerful reductant, requires anoxic conditions. Accordingly, the likely acquired peroxidases include cytochrome c peroxidase (Ccp), which could scavenge hydrogen peroxide in the periplasm [77], and the most prevalent peroxidase [78], thioredoxin-dependent peroxiredoxin (AhpC). Besides peroxide scavengers, the desulfoferrodoxin (Dfx), which functions as superoxide reductase (SOR) to reduce superoxide to hydrogen peroxide and which is broadly distributed among anaerobic bacteria [78], were likely acquired during the origin of anammox bacteria (Fig. 3). Another acquired gene is *fprA*, which encodes flavo-diiron proteins that scavenge O_2_.

Investigating other metabolisms that were also acquired upon the origin of anammox bacteria allows reconstructing the coeval ecology. Our data show that *sat* (Fig. 3), which encodes sulfate adenylyltransferase for assimilatory incorporation of sulfate into bioavailable adenylyl sulfate, as well as *aprAB* and *fsr* (Fig. 3) for encoding dissimilatory sulfite reductase, were acquired upon the origin of anammox bacteria (Fig. 3). This acquired capability of sulfate metabolism is coincident with the increased concentrations of seawater sulfate from ∼100 µM or less throughout much of the Archean [79, 80] to over 1mM after the rise of atmospheric O_2_ [81, 82]. Unlike other Planctomycetes, anammox bacteria are generally autotrophs that use the Wood-Ljungdahl pathway for carbon assimilation [83]. Consequently, they potentially lost many genes involved in carbohydrate utilization (Fig. 3). The metabolic loss is further strengthened by the enrichment analysis where the genes predicted to be lost by the LCA of anammox bacteria are enriched in pathways involving hydrolase activity, intramolecular oxidoreductase activity and carbohydrate kinase activity (Fig. 3; see Data and Code availability), hinting at a decreased ability of anammox bacteria to degrade organics.

Additionally, iron is a vital element for HZS [9] and HDH [84], and *in vitro* studies revealed that increased iron concentrations can promote the growth of anammox bacteria [85]. The ancient oceans are thought to have been ferruginous (high iron availability) according to sedimentary records of iron speciation across several coeval marine settings from the Archean through most of the Proterozoic [86], meaning that large amounts of soluble iron would have been available for anammox bacteria. Interestingly, a series of iron-related genes to make use of the abundant iron from the environment were likely acquired upon the origin of anammox bacteria. First, anammox bacteria likely acquired the gene *fur*, which encodes a ferric uptake regulator for iron assimilation. Moreover, the iron-containing proteins, specifically cytochrome, are encoded by the acquired genes *petBC*, which potentially encodes Rieske-heme b [13], and *CcsAB*, which encodes a protein involved in cytochrome c maturation systems (Fig. 3). Notably, *ahbABCD*, which encodes proteins-synthesizing *b*-type hemes in anammox bacteria [87], were likely acquired before the origin of anammox bacteria (Fig. 3). Furthermore, we identified that the gene *Bfr* that encodes the oligomeric protein, bacterioferritin involved in the uptake and storage of iron [88] was likely acquired during the origin of anammox bacteria (Fig. 3), which may help to hoard iron in settings where iron became locally sparse, such as along euxinic (sulfide-rich) or oxic marine margins in the Paleoproterozoic [89]. A recent study highlights the vital role of bacterioferritin to the anammox regulation [90]. It is thought that the appearance of euxinic margins around the time of the GOE trapped iron that upwelled from the deep ocean, leading to the demise of iron oxide deposits (i.e., banded iron formations) [91]. Our results tentatively support this model, if the microenvironments that those primordial anammox bacteria colonized were located near the euxinic-oxic interface where nitrite and ammonium were available while upwelled iron was titrated out of the water column by freely dissolved hydrogen sulfide.

Taken together, the timelines and corresponding genomic changes of anammox bacteria provide implications for the physiological characteristics of descendant anammox bacteria and the origin of other nitrogen-transforming pathways in the context of an estimated evolutionary timeline of anammox bacteria. For example, the earlier branching lineage *Ca.* Scalindua exhibit high nitrite affinity (0.45 μM) compared to other anammox genera (up to 370 μM) [15, 92], which may reflect a progressive growth of the marine nitrite reservoir, linked to the protracted oxygenation of the surface environments over geologic timescales [29]. Likewise, genomic data reveal that also canonical denitrification radiated across the tree of life after the GOE [93], and the gains (Fig. S10; see Supplementary Text section 4.4) of genes *nrfAH* for dissimilatory nitrate reduction to ammonium (DNRA) at the late-branching lineages (*Ca.* Brocadia and *Ca.* Kuenenia) at around 800 Ma are consistent with the time when the nitrite/nitrate availability was increased by the Neoproterozoic Oxygenation Event [94]. The time estimate and genomic changes of the anammox bacteria have gone some way towards enhancing the understanding of the physiological characteristics of modern anammox bacteria and the historical N cycle.

## Caveats and concluding remarks

In the present study, we link an estimated evolutionary timeline of anammox and geological context of early Earth on a simplified view that nitrite-dependent anammox is ancestral to anammox bacteria. However, there are several limitations. First, dating the bacterial evolution has many challenges, which, among others, include the paucity of appropriate fossils, and a long evolutionary distance between anammox bacteria and cyanobacteria [51]. Hence, the date estimates should be read as potential ranges of time (indicated by the 95% HPD interval), instead of accurate time points, under the molecular clock model. Moreover, it is worth pointing out the metabolism called feammox, a process where ferric instead of nitrite is taken as the electron acceptor for ammonia oxidation. Both feammox and anammox can anaerobically convert ammonia into dinitrogen [95], but the former process is supposed to be carried out by Acidimicrobiaceae from the Actinobacteria phylum [96, 97] and under anaerobic environments without the presence of anammox bacteria [98]. Likewise, a recent experimental study [99] proposed extant anammox bacteria as electroactive organisms and suggested the feasibility to utilize graphene oxides and man-made metallic electrodes mimicking metal oxides as the electron acceptor. However, the lack of detailed molecular mechanism hampers the inference of its evolutionary history. In any case, metal oxides such as Fe_2_O_3_ or MnO_2_ also became more abundant in the Neoarchean to Paleoproterozoic, i.e. around the time of the GOE [91, 100], and hence even if these oxides were used as substrates for anammox, it would not violate our overall conclusion that the GOE played a primary role in driving the evolution of this metabolic pathway. However, it is also worth noting that metal oxides are solids and thus require extra-cellular electron transfer [99]. An entirely intra-cellular metabolism, using dissolved nitrite, may be more likely to evolve first, especially given that nitrate became bioavailable in the ocean around the same time (see above).

Another important aspect to highlight about our approach is that, our analysis began with inferring the LCA of anammox bacteria based on the presence of *hzsCBA*, a common strategy in phylogenetic studies. It is important to note that such an enzyme complex encoded by multiple genes might originate in different genomic backgrounds and were presented together in a suitable genomic background later in evolution. This scenario, if true, indicates that the appearance of any single subunit of *hzsCBA* genes, which could not assembly a functional hydrazine synthase, should predate the LCA of anammox bacteria. Nevertheless, the alternative hypothesis would not affect our inference and following analysis since we only focus on the inferred ancestor of extant anammox bacteria, a strategy commonly used in modern phylogenetic analysis. These alternative possibilities, although not affecting the molecular clock analysis, should be carefully considered when making inference based on our estimated evolutionary timeline of anammox bacteria.

Here, using molecular dating and comparative genomics approaches, we link the emergence of an obligate anaerobic bacterial group, which drives the loss of fixed N in many environments, to the rise of O_2_, and highlight their evolutionary responses to major environmental disturbances. Apparently, the GOE opened novel niches for the origin and subsequent expansion of diverse aerobic prokaryotes such as AOA and AOB, which in turn facilitated the origin of other bacterial lineages, including some anaerobes like the anammox bacteria, by providing the resources for their energy conservation. Our results open up the possibility that a significant proportion of the Proterozoic nitrite budget was consumed by anammox bacteria, and that the sedimentary nitrogen isotope record might be influenced by their activity. Our findings may have implications for Proterozoic climate, because N_2_O has previously been invoked as an important greenhouse gas at that time [101], but a relatively higher contribution of anammox bacteria to the marine nitrogen cycle may have hampered N_2_O production [6]. Lastly, our study suggests that the impacts of the GOE go well beyond aerobic prokaryotes. We therefore conclude that molecular dating is a likely feasible approach to complement isotopic evidence for resolving the timeline of biological evolution and for providing additional constraints on climate models of the distant past.

## Materials and Methods

Overall, we compiled two genome sets and one 16S rRNA gene set in our study (Dataset S1.1). A phylogenomic tree (Fig. S1) was generated based on the concatenated alignment of 120 ubiquitous proteins (bac120; Dataset S1.3) proposed for tree inference by the Genome Taxonomy Database [102]. All published genomic sequences (952 in total) affiliated with the phylum Planctomycetes in NCBI Genbank up to December 2019 were retrieved (Dataset S1.2). To further examine whether the patterns obtained with this Planctomycetes dataset hold the same, we further constructed an expanded set of Planctomycetes genomes by retrieving a total of 2,077 Planctomycetes genomes released by April 2021 at Genbank (Fig. S2; see Data and Code availability). Key anammox genes *hzsCBA* and *hdh* were identified against manually curated reference sequences by BLASTP. For each gene, identified protein sequences were aligned using MAFFT (v7.222) [103] and the alignments were refined by trimAl (v1.4) [104]. All phylogenies were constructed by IQ-tree (v1.6.2) [105] with substitution models automatically selected by ModelFinder [106] and branch support assessed with 1,000 ultrafast bootstrap replicates. Note that the constraint topologies for dating analysis (Fig. 1; Fig. S5) were inferred with the profile mixture model (LG+C60+G) which better accommodates across-site heterogeneity in deep-time evolution. Following manual curation, four well-recognized genera and two separate lineages of anammox bacteria were highlighted with different colours, and the presence of key genes were annotated with filled symbols beside the labels (Fig. S1). Furthermore, the phylogenomic tree was generated with similar methods using 16S rRNA genes identified from downloaded genomes and those retrieved from the SILVA database (Fig. S3; Dataset S1.3). All trees (including phylogenomic and 16S trees) in our study were visualized with iTOL [107]. More details are shown in supplementary text, section 2.

Molecular dating analysis was carried out using the program MCMCTree from the PAML package (4.9j) [108]. In our study, 13 calibration sets constructed with different time constraints of the root and three calibration nodes within the cyanobacteria lineage were used (see Supplemental Texts section 3.2). The topology constraint for dating analysis was generated using 85 genomes, which were sampled from the phylogenomic tree of phylum Planctomycetes (see Supplemental Texts section 3.1), and the model LG+C20+F+G under posterior mean site frequency (PMSF) approximation [109]. To perform dating analysis, clock models and different calibration sets were compared in 26 schemes (see Supplemental Texts section 3; Dataset S2.1) [110, 111]. For each scheme, the approximate likelihood method [112] of MCMCTree were conducted in duplicate with identical iteration parameters (burn-in: 10,000; sample frequency: 20; number of sample: 20,000). The convergence of each scheme was evaluated by comparing the posterior dates of two independent runs (see Data and Code availability). With the updated genome set 2 (Dataset S1.1), we repeated the taxon sampling process and generated an alternative phylogeny for dating analysis (Fig. S5) using LG+C20+F+G model (C20: 20 classes of site-specific amino acid profiles) under PMSF approximations. Similarly, we also generated another phylogeny for dating analysis using the same genome set as used in Fig. 1 but with the profile mixture model (LG+C60+G) which uses 60 classes of amino acid profiles [113]. Note that the missing parameter +F would not significantly change the results since it only adds another profile amino acid calculated from the original data. These two repeat dating analyses were conducted with the best practice dating scheme (IR model and calibration C1). Moreover, we performed likelihood based dating analyses [114] (Fig. S9) using the phylogenomic tree (Fig. S2) and the calibration C1 (see Data and Code availability) implemented in ape [56].

The phylogenetic differences between gene trees and the species tree were reconciliated by GeneRax [115] with unrooted gene tree as input and automatically optimized duplication, transfer, loss (DTL) rates. For each gene identified from the genome set 2 (Dataset S1.1), we used a species tree comprised by all anammox bacteria pruned from the phylogenomic tree (Fig. S2) as the reference. We used recommended parameters including SPR strategy, undated DTL model and a maximum radius of five.

For comparative genomics analysis, the protein-coding sequences were annotated against KEGG [116], CDD [117], InterPro [118], Pfam [119], TIGRFAM [120] and TCDB [121], individually (see Supplemental Texts section 4). Following annotations, the potentially gained/lost genes between groups (anammox bacteria versus non-anammox bacteria; an anammox bacterial genus versus all other anammox bacteria) were statistically tested using two-sided fisher’s exact tests. Finally, the resulting *p*-values were corrected with the Benjamini-Hochberg FDR (false discovery rate) procedure. Moreover, genes with corrected *p*-values smaller than 0.05 and with larger ratio in the study group were defined as gained genes, and genes with corrected *p-*values smaller than 0.05 and with smaller ratio in the study group were defined as potentially lost genes. Those potentially gained/lost genes were summarized for their detailed functions (see Data and Code availability).

## Acknowledgements

We thank Bite Pei for building the basic dataset for this project, Yang Qian for his help in editing the earlier version of the manuscript, Jinjin Tao for her helpful discussion, and Mario dos Reis for his suggestion on dating analysis. This work is funded by the National Science Foundation of China (92051113), the Hong Kong Research Grants Council Area of Excellence Scheme (AoE/M-403/16), the Direct Grant of CUHK (4053495), and The CUHK Impact Postdoctoral Fellowship Scheme to (S.W.).

## Data and Code availability

All used genomic sequences, generated phylogenetic trees, estimated divergence times and the used python codes are deposited in the online repository https://github.com/luolab-cuhk/anammox-origin.

