## Supplementary information for "Phylogenomic evidence for the Origin of Obligately Anaerobic Anammox Bacteria around the Great Oxidation Event"

**This PDF file includes:**

Supplementary Text

Figures S1 to S9

References

**Supplementary Text**

**1 Genome retrieval and gene predictions**

*1.1 Genome retrieval*

*1.2 Gene prediction*

**2 Phylogenetic analysis**

*2.1 Phylogenomic tree based on bac120 proteins*

*2.2 Phylogenomic tree based on 16S ribosomal RNA (rRNA) genes*

*2.3 Phylogeny of key genes for anaerobic ammonia oxidation (anammox)*

**3 Molecular dating analysis of Anammox bacteria**

*3.1 Taxon sampling and topology constraints for dating analysis*

*3.2 Justification of calibrations*

*3.3 The effect of the clock models*

*3.4 The effect of calibration constraints*

*3.5 Time estimates using mitochondria-based strategy*

**4 Comparative genomics analysis**

*4.1 Annotations against six databases*

*4.2 Fisher's exact tests*

*4.3 Enrichment analysis*

*4.4 Genomic changes between genera within anammox bacteria*

### 1. Genome retrieval and gene predictions

#### *1.1 Genome retrieval*

We retrieved 952 genomic sequences of Planctomycetes including anammox bacteria from GenBank of NCBI on December 02, 2019 (Datasets S1.1 & S1.2). Retrieved genomic sequences comprise single amplified genomes (SAGs), metagenome-assembled genomes (MAGs) and whole-genome sequencing (WGS) of enriched culture samples (a few anammox bacteria) or isolates. Six Verrucomicrobia genome sequences were also downloaded to serve as outgroup species. To estimate the age of anammox bacteria, bacterial lineages with fossil calibrations need to be included in a molecular dating analysis. Since the lineage-informative bacterial fossils are only found in the phylum Cyanobacteria, 21 high-quality reference genomic sequences of oxygenic Cyanobacteria and 10 genomic sequences of Melainabacteria were downloaded from RefSeq and GenBank. The detailed environmental information of each genomic sequence was retrieved through Entrez Kans J [1] of NCBI using custom scripts (Dataset S1.2; see also Data and code availability). All downloaded genomic sequences were re-annotated with Prokka (v1.14.5) [2]. The completeness of downloaded genomes was estimated with taxonomic-specific workflow (using corresponding phylum) of checkM (v1.1.1) [3]. As the size of genomic data sets expands rapidly in recent years, an updated dataset comprising 2,077 Planctomycetes genomes was retrieved from GenBank (April 20, 2021) to confirm the phylogenetic structure and the age of anammox bacteria. The usage of datasets and related figures/analyses is briefly summarized in the Dataset S1.1.

#### *1.2 Gene prediction*

The predicted proteins were annotated by hmmer (v3.2.1) [4] against pre-built HMM profiles of the 120 bacterial proteins (bac120) which have been widely used in tree inference for bacteria [5], with an E-value cutoff of 1e-50 for phylogenetic analysis (Dataset S1.4). To perform molecular dating analysis, the predicted proteins were annotated by hmmer against the profiles of 25 universally conserved genes across bacteria (cog25) [6] with an E-value cutoff of 1e-20 (Dataset S1.4). We filtered out genomes with less than 24 (20%) genes annotated by the bac120 proteins, and 881 genomes were retained for subsequent analysis (Dataset S1.2). Hence, 887 genomes (881 sampled Planctomycetes and six high quality Verrucomicrobia) were used in phylogenetic and dating analyses.

#### 2. Phylogenetic analysis

##### *2.1 Phylogenomic reconstruction based on bac120 proteins*

The annotated proteins against bac120 were aligned at amino acid levels using the G-INS-I refinement method of MAFFT (v7.222) [7], and spurious sequences or poorly aligned regions were trimmed using trimal (v1.4) [8] with recommended parameters (resoverlap: 0.55; seqoverlap: 60). The maximum likelihood (ML) phylogenomic tree comprising 881 Planctomycetes genomes and six Verrucomicrobia genomes taken as the outgroup was constructed by IQ-Tree (v1.6.2) [9]. The ML phylogenomic tree (Fig S1) was built based on the concatenated alignment of the retrieved bac120 proteins from each genome, with the parameters '-mset WAG,LG,JTT,Dayhoff -mrate E,I,G,I+G -mfreq FU' and 1,000 replicates of ultrafast bootstrap [10]. 'LG+I+G4' was chosen as the best-fit model after model tests by

ModelFinder [11] implemented in IQ-Tree. Note that phylogenomic reconstruction based on profile mixture model, which likely better accommodates across-site heterogeneity in the amino acid substitution [12], is computationally infeasible due to the large number of analyzed genomes (n=887). Instead, we used profile mixture model (LG+C20+F+G) to infer a phylogeny for only molecular clock analysis with a reduced genome set consisting of 85 taxa (see Section 3). To examine the robustness of the patterns obtained with this Planctomycetes dataset, we further constructed an expanded set of Planctomycetes genomes by retrieving a total of 2,077 Planctomycetes genomes (Genome set 2; see Dataset S1.1) released in April 2021 at Genbank (Fig. S2; see Data and Code availability) with the model best-fit substitution model LG+I+G4. All trees (including phylogenomic and 16S trees) in our study were visualized with iTOL v5 [13].

Two lineages of anammox bacteria were newly identified in our study, namely the 'basal lineage' and the '*hzsCBA*-less lineage' (Figure S1), both of which were represented by the MAGs collected from the underground aquifer system adjacent to the Colorado River [14]. The study employed three different field experiments including time-series samples across the duration of acetate amendment, time-series samples across the duration of oxygen injection, and time-series samples from natural high- (3.602 mM) and low-oxygen (0.287 mM) conditions in the groundwater, driven by fluctuations in the water table in situ. The 'basal lineage' named in our study came from time-series samples with varying oxygen conditions, while the '*hzsCBA*-less lineage' came from field experiments after acetate or oxygen injections. Moreover, both lineages were exclusively observed in samples collected with 0.2 μm filter, and consequently, they likely adapt free-living lifestyles which are rarely observed in most anammox bacteria [15].

*2.2 Phylogenetic tree based on 16S rRNA genes*

We retrieved 20,142 rRNA genes of the class Brocadiae comprising both anammox and non-anammox bacteria within Planctomycetes from SILVA (r138) [16], and further clustered them into 913 clusters with cd-hit v4.8.1 [17] using 0.97 [18] as an identity threshold for thoroughly investigating the phylogenetic relationship of anammox bacteria. Since properly selected reference sequences could help us identify lineage affiliations of the sequences derived from the uncultivated organisms, we identified 16S rRNA genes in 457 out of the 958 genomes affiliated with Planctomycetes and the outgroup Verrucomicrobia (Dataset S1.3) as references. We further incorporated 16S rRNA genes from genomic sequences used in the phylogenetic inference: i) 42 16S rRNA genes from anammox bacteria genomes for identifying each anammox genus, ii) three 16 rRNA genes from non-anammox Planctomycetes genomes (GCA_003567495.1, GCA_003551305.1, GCA_001303885.1) which belong to the sister lineage to anammox bacteria to assist identification of anammox bacteria, and iii) six 16 rRNA genes from the Verrucomicrobia genomes to help identify Planctomycetes genomes. Together with the representative sequences of generated 913 clusters, the 16S rRNA gene tree (Fig. S3) was generated using the same parameters described above.

Consistent with a previous study [19], all anammox bacteria used in our study show a monophyletic origin in both phylogenies using the bac120 marker genes and the 16S rRNA genes. Note that the anammox bacterium (*Ca.* Brocadia *sp.* SB37) showed different phylogenetic positions between the phylogenomic tree (Fig. S1) and the 16S rRNA gene phylogeny (Fig. S3). Furthermore, the genus *Ca.* Anammoximicrobium which was reported to be a presumable anammox bacteria lineage based on its 16S rRNA gene KC467065.1 [20], was not included in our study. This is because the latest genomic sequence (GCA_012515235.1) from this genus [21] which displays a high 16S rRNA gene sequence similarity (96.88%) to the first reported 16S rRNA gene (KC467065.1), does not contain any anammox genes (see Data and code availability) despite a high completeness (96.74%) of the genome. This calls for careful validations of the anammox ability of this genus.

##### *2.3 Phylogenies of key genes for anammox*

The phylogenetic analysis of metabolic genes could provide direct insights into the evolution of metabolism. Anammox relies on enzymes encoded by *hdh* and *hzsCBA* [22]. The gene *hdh* encoding hydrazine dehydrogenase for anammox was not included in phylogenetic analysis because of its extensive duplications [23], which could confound phylogenetic analysis. The predicted proteins from 2,077 Planctomycetes genomes were annotated using BLAST with an E-value cutoff of 1e-20 against manually curated HzsCBA proteins. Only *hzsCBA* genes within the same gene cluster were used to build the gene tree (see Data and Code availability). Amino acids of these identified HzsCBA were aligned with MAFFT (v7.471) [7], and the generated alignments were trimmed with trimAl (v1.4) [8]. Gene phylogenies were built using IQ-TREE (v1.6.2) [9] with the key parameters '-m TEST -madd LG+C20+G,LG+C30+G,LG+C40+G,LG+C50+G, LG+C60+G -mset WAG,LG,JTT -mrate E,I,G,I+G' which allow the best-fit model selection to range from amino acid exchange rate matrices and profile mixture models C20-C60. All three phylogenies were inferred with the same best-fit model (LG+C60+G). As far as we know, no available outgroups of *hzsCBA* were reported. We therefore turned to the minimal ancestor deviation (MAD) [24] to infer the root of the phylogeny of each gene (Fig. S4). This rooting method accommodates lineage variation in evolutionary rates by using all pairwise metric and topological information, outcompeting existing rooting methods which might fail to accommodate the evolutionary rate heterogeneity. The rooted trees were further reconciliated with the species tree by GeneRax (v1.2.2) [25]. The software implements a species-tree-aware approach which unites inferences of gene phylogeny and HGT events according to established maximum likelihood optimization algorithms. The reconciliation results suggest a single origin of *hzsCBA* from the LCA of anammox bacteria (Fig. S4), confirming the results shown by the phylogenomic tree and the 16S rRNA gene tree.

The phylogenetic differences between gene trees and the species tree were reconciliated by GeneRax (v1.2.2) (Morel et al. 2020) with unrooted gene tree as input and automatically optimized duplication, transfer, loss (DTL) rates. For each gene, we used a species tree comprised by all anammox bacteria pruned from the phylogenomic tree (Fig. S2) as the reference. We used recommended parameters including SPR strategy, undated DTL model and a maximum radius of five.

#### 3. Molecular dating analysis of Anammox bacteria

Multiple parameters employed in dating analysis would significantly affect the posterior time estimate. To perform a careful molecular dating analysis, the divergence times of anammox bacteria were estimated using 26 different schemes (Dataset S2.1). These schemes differ in the choice of calibration constraints and clock models [auto-correlated rates (AR) and independent rates (IR) model], which are often the two factors that mostly influence dating based on prior studies [26]. We used the 25 universally conserved genes (Dataset S1.4) termed as cog25 [6] as the sequences in dating analysis. For each scheme, two repeats were run separately with the identical parameters (burn-in: 10,000; sample frequency: 20; number of samples: 20,000). We ensured that convergence had reached by comparing the estimated parameters from two independent runs (see Data and Code availability). The repeated analysis using the latest Planctomycetes genomes including a total of 2,077 Planctomycetes genomes released in April 2021 at Genbank, was conducted with the best practice dating strategy (IR model and calibration C1). Also, to avoid potential impact brought by the constraint topology, we built the topology with LG+C60+G model (see section 3.1) and repeated the dating analysis with the best-practiced strategy. In general, the above two repeat analyses yielded similar time estimates of the LCA of the anammox lineage (Fig. S6) and the related results are deposited at the online repository (see Data and Code availability).

*3.1 Taxon sampling and topology constraints for dating analysis*

To reduce computational costs of Bayesian estimation in divergence times, taxon sampling was applied considering following criteria: 1) Sampling the basal lineage according to the anammox bacteria in this study, and 2) Sampling the genomes with more marker genes for dating. Accordingly, we selected 85 genomes, where the Melainabacteria group was used as the outgroup of oxygenic Cyanobacteria [27], from the full set of 887 genomes (see Supplementary Text section 1.2) with TreeCluster v1.0 using Threshold-Free approach [28]. This ensures that among all of the 25 conserved genes used in molecular clock analysis (20040 sites after alignments and concatenation), there are four backbone genes (those shared by all analyzed genomes) which should be sufficient to provide robust date estimates according to a prior study [29]. The phylogenomic tree with the concatenated alignment of annotated bac120 proteins from 85 genomes including 48 genomes of Planctomycetes (including 37 anammox bacteria), six Verrucomicrobia, 21 oxygenic Cyanobacteria and 10 Melainabacteria genomes was built using IQ-Tree (v1.6.2). The model LG+C20+F+G under posterior mean site frequency (PMSF) approximation [30] was applied based on the alignments of the concatenation of bac120 proteins (Dataset S1.4) [5]. The support values were calculated with 1,000 replicates using the ultrafast bootstrap algorithm [10]. The topology of the generated tree is highly similar to the aforementioned full phylogeny involving 887 Planctomycetes genomes. The Melainabacteria group and Cyanobacteria were used to root the tree. The placement of calibration nodes and time constraints were shown in Figure 1 and Dataset S2.1.

##### *3.2 Justification of calibrations*

The molecular dating analysis heavily depends on the use of time calibrations. In the case of bacterial tree of life, calibrations are only available in Cyanobacteria. In our study, we used four calibrations, which constrain the root, the origin of oxygenic Cyanobacteria, the origin of *Nostocales* and the origin of *Pleurocapsales*. Here is a summary of the calibrations we used.

For the root, which is the LCA of Planctomycetes and Cyanobacteria/ Melainabacteria, the minimum age cannot be determined due to the lack of fossils. Thus, we only adapted the maximum age according to the earliest evidence of life on Earth. It is proposed that life likely emerged after the late heavy impact at 3,800 million years ago (Ma) [31]. However, the timing of the impact has been debated and varies according to different biogeochemical evidence [32]. Others suggested that some organisms might have survived the impact [33], thus supporting an older origin of life. Accommodating the above uncertainties, the largest time estimate of the impact [34] which coincides with the age of the Earth (4,500 Ma) [35] was set as the upper bound of the root. On the other hand, the stromatolites of Pilbara Supergroup (3,490 - 3,240 Ma) provide the oldest convincing evidence of life on Earth [36]. Accordingly, we also employed 3,500 Myr as the maximum bound of the root age.

The time constraints for the origin of oxygenic Cyanobacteria are heavily debated. According to Sanchez-Baracaldo P, Raven JA, Pisani DKnoll AH [37], two maximum ages based on the rise of atmospheric oxygen at 3,000 Ma [38] and the presence of hopanes at 2,700 Ma [39], and one minimum age which is based on the Great Oxidation Event (GOE) ~2,300 Ma, were implemented. However, as critically discussed in Zhang H, Sun Y, Zeng Q, Crowe SALuo H [40], the use of the maximum time constraint in these calibrations in Sanchez-Baracaldo P, Raven JA, Pisani DKnoll AH [37] is not appropriate: the presence of evidence for the rise of $O_2$ should not be used as a maximum age, but rather, it suggests that oxygenic Cyanobacteria could have appeared before GOE ~2,300 Ma. Furthermore, the sedimentary records of chromium isotopes and redox-sensitive metals suggest that atmospheric oxygen had reached an appreciable level by 3,000 Ma [38], which indicates an earlier origin time of oxygenic Cyanobacteria. Overall, we set either 2,300 Myr or 3,000 Myr as the minimum bound of the total group of oxygenic Cyanobacteria and no maximum bound was set.

The time constraints for the total group of *Nostocales* or *Pleurocapsales* are also contentious. Since the upper bounds of those are hard to resolve, we followed Zhang H, Sun Y, Zeng Q, Crowe SALuo H [40] to set 1,700 Myr as the minimum bound of *Pleurocapsales* based on the presence of microfossils [41], and set 1,600 Myr as the minimum bound of *Nostocales* based on the discovery of the nostocalean akinetes fossil [42], which are high-confidence fossils that represent these two cyanobacteria orders, respectively.

##### *3.3 The effect of the clock models*

The auto-correlated rates (AR) model proposes that evolutionary rates of descendants are correlated with their parental rates, whereas the independent rates (IR) model assumes independent rates among lineages [43]. Different molecular clock models (IR vs. AR) resulted in 1~12% difference in time estimates (Dataset S2.1). We used the stepping stone method with the approximate likelihood to estimate the marginal likelihood of different molecular clock models [44-46]. We selected calibration sets C7, C13, C16 and C21 which differs at the root or the total group of oxygenic Cyanobacteria to be shown in Figure 1. Each scheme was sampled from eight repeats with different powers generated using mcmc3r [43]. The log marginal likelihood of each model and the corresponding Bayes Factor (BF) between the two competing models were summarized in Dataset S2.2. All BFs using different schemes favored the IR model, which was therefore used as the molecular clock model in the main analysis.

##### *3.4 The effect of calibration constraints*

The geological evidence which defines the boundaries of some calibration constraints used here remains debated in their biogenicity and interpretation [47]. Likewise, the biological affinities of early fossils may be controversial [48]. Thus, the calibration constraint used in molecular dating analysis should be carefully benchmarked.

We selected calibration C1 as the best-practiced calibration set for the following reasons. First, analysis using it displayed the highest precision, as reflected by the lowest slope in the infinite-sites plot (Fig. S6), where the widths of 95% highest posterior density

(HPD) interval is plotted against the posterior mean ages of nodes [49]. Second, calibration C1 showed a relatively similar overlap in the LCA of anammox bacteria between the prior and posterior time distributions among other calibration sets (Fig. S7), suggesting that its calibrations used are as informative as others [50].

##### *3.5 Time estimates using mitochondria-based strategy*

Apart from the molecular dating analysis based on cyanobacterial calibrations, a novel molecular dating strategy based on the mitochondrial endosymbiosis event has recently been developed [26]. To perform mitochondria-based dating analysis, we compiled two gene sets. The first set was based on the 24 mitochondria-encoded genes used in the original study [26] (mito24: 6295 amino acids). Because some of these genes, which were originally identified to be shared by mitochondria and (α-)proteobacteria, might not have clear orthologs in Planctomycetes, we additionally selected six genes (mito6: 1238 amino acids) according to the following two criteria; i) these genes should be found in over 85% of genomes used to construct the phylogeny of the bacterial tree of life in a recent phylogenomics study [51], and ii) they are included in at least one of the following three sets of genes conserved across the bacterial tree of life: bac120 [5], Coleman2021 [51], and cog25 [6]. The list of genes is provided in Dataset S1.4.

The genomes analyzed were sampled from Genome set2 (Dataset S1.1), and the topology was generated using the LG+C60+G model (see section 3.1). According to recent phylogenomic studies, the mitochondria lineage was placed as the sister lineage to αproteobacteria [26, 52, 53], and they formed together a monophyly placed as a sister lineage to the PVC group[54, 55], which is a superphylum of bacteria that includes Planctomycetes, Verrucomicrobia, and Chlamydiae (Fig. 2A).

Apart from the three cyanobacterial calibrations (Node 2 to Node 4), four eukaryotic calibrations (Node 5 to Node 8) were implemented and shown in the Fig. 2, according to [26, 56]. These calibrations corresponded to the total group of Bangiophyceae (crown group of red algae) (minimum: 1033 Ma), the total group of florideophytes (minimum: 550 Ma), the total group of mosses (crown group of Embryophyta) (maximum: 509 Ma & minimum: 450 Ma), and the total group of eudicots (crown group of angiosperms) (maximum: 250 Ma & minimum: 125 Ma). More details of time constraints are provided at Dataset S2.1 (see also the legend of Figure 2).

#### 4. Comparative genomics analysis

##### *4.1 Annotations against six databases*

To comprehensively investigate potential gene gains and losses, the gene annotations based on six different databases including KEGG [57], CDD [58], InterPro [59], Pfam [60], TIGRFAM [61] and TCDB [62] were searched against with an E-value cutoff of 1e-20.

Among the 3,096,667 coding sequences in the 887 genomes, nearly 60% of them were successfully assigned to a functional category in at least one of the six databases. Particularly, the KEGG database annotated nearly 50% of all coding sequences, which is the most among all databases.

*4.2 Fisher's exact tests*

We individually performed two-sided fisher's exact test against a binary (presence or absence) table of each annotated functional category. The derived *p*-values were corrected using the Benjamini-Hochberg FDR (false discovery rate) procedure. The comparisons of anammox bacteria versus non-anammox bacteria (those within a blue box in Fig. 3) and between different genera within anammox bacteria were performed. Genes with corrected *p*-values smaller than 0.05 and with smaller ratio in the study group (anammox bacteria) than in the reference group (non-anammox bacteria) were defined as potentially lost genes in anammox bacteria. The presence or absence of functional categories which are ecologically important was visualized against each genome along with the phylogenomic tree using iTOL v5 (Figs. 3 and S9) [13]. Besides, for gene annotations of the 34 ladderane synthesis candidate genes, we downloaded the protein sequences from Rattray JE, Strous M, Op den Camp HJ, Schouten S, Jetten MSDamste JS [63] and searched for homologs in the analyzed genomes by BLAST with an E-value cutoff of 1e-20. The detailed information of the presence or absence of these 34 genes among all Planctomycetes genomes is provided at github (see Data and code availability).

Although the Fisher's exact test-based approach unveiled the significantly different genes based on the presence or absence table between anammox and non-anammox bacteria, (Fig. 3), it should be kept in mind that this is a simple model without considering branch lengths, phylogenetic incongruence and genome incompleteness. Also, the detailed functions of many presumably gained/lost genes identified by this approach remain further investigation (see Data and code availability).

*4.3 Enrichment analysis*

The GOATOOLS (v1.0.3) [64] was used to perform enrichment analysis against acquired and lost genes, separately. Since the KEGG database annotated the most genes against other five databases, only annotations from KEGG database were used to perform enrichment analysis in our study. To acquire the hierarchy information, the KEGG annotations were converted into IPR annotations through Interpro database. The enrichment analysis requires two files with IPR IDs: one file is a population file containing the list of the genes (represented by their IPR IDs) present in all 65 anammox bacteria genomes taken as background, and the other is a study file containing the list of genes present in the target group. The enrichment results are provided in github (see Data and code availability).

*4.4 Genomic changes between genera within anammox bacteria*

Our phylogenomic analysis revealed six clades within anammox bacteria, which correspond to four known genera of anammox bacteria (*Ca.* Brocadia, *Ca.* Jettenia, *Ca.* Kuenenia, *Ca.* Scalindua), and two lineages newly defined in the present study (the basal lineage and *hzsCBA*-less lineage) (Fig. 1). To gain insight into the differences in the genome content between different lineages, the presence or absence of identified genes in each lineage were individually compared with the genomes of the other anammox bacteria lineages with multiple fisher's exact tests.

The anammox bacteria are usually classified into two physiological groups including freshwater anammox (*Ca.* Brocadia, *Ca.* Jettenia and *Ca.* Kuenenia) and marine anammox (*Ca.* Scalindua) [65]. The presence of sodium-proton antiporter might be responsible for the salinity adaptation of *Ca.* Scalindua, the dominant anammox bacteria lineage in modern marine environments [66, 67]. However, the gene *rnf* encoding NADH hydrogenase for sodium translocation, which plays a role in adapting to the high-salinity environment based on experimental studies [68], was plausibly lost in marine anammox bacteria *Ca.* Scalindua (Figs. 3 and S9). Interestingly, Huang XW, Mi WK, Ito HKawagoshi Y [69] has studied the abilities of salinity resistance among different genera of anammox bacteria and found that even some freshwater species displayed high salinity resistance, which we predict might be attributed to the presence of *rnf* in their genomes. Besides, the gene *htpG* encoding a heat shock protein, appears almost specific to *Ca.* Scalindua (Fig. S9), which likely provides thermotolerance to *Ca.* Scalindua in deep-sea hydrothermal vents and geothermal subterranean oil reservoirs [70, 71]. Intriguingly, the genes *hdrABCD* encoding heterodisulfide reductase, which catalyzes disulfide reduction assisted by iron-sulfur cluster [72], were found common in *Ca.* Scalindua but not in other anammox bacteria lineages, but their detailed roles need further investigation. Genes lost in this lineage included *macB*, *fadL*, *AraJ*, and *msmX* (Fig. S9) which are likely responsible for exportation of macrolide, fatty acid, multidrug and multiple sugars, respectively, suggesting less extracellular organics produced by *Ca.* Scalindua in anammox consortia [73].

The other three anammox genera, *Ca.* Brocadia, *Ca.* Jettenia and *Ca.* Kuenenia show specific metabolic patterns which might help to understand their physiological differences. The acquired genes *mexLJK* of *Ca.* Kuenenia (Fig. S9) involved in multidrug efflux system which might explain its tolerance to penicillin G and chloramphenicol reported in previous studies [74, 75]. Further, the permease proteins encoded by *Sulp* and *Ycjp* (Fig. S9) for the acquisition of sulfur and different sugars might provide metabolic versatility to *Ca.* Kuenenia [76]. The presence of gene *hmp* (Fig. S9) encoding nitric oxide dioxygenase hints its unusual ability to oxidize nitric oxide. The key gene *nrfA* encoding pentahaem nitrite reductase for dissimilatory nitrite reduction to ammonia is absent in the genomes of all *Ca.* Kuenenia (Fig. S9). Instead, an unusual calcium-dependent trihaem nitrite reductase encoded by *CaNIR* (Fig. S9) has been reported in *Ca.* Kuenenia bacteria [77]. The ubiquity of *CaNIR* in three freshwater genera but absence in *Ca.* Scalindua implies its functions during habitat expansions from marine to freshwater. Further, the likely acquired *fetAB* (Fig. S9) encoding iron exporter protein might provide the ability to protect against oxidative species through exporting iron [78].

All analyzed *Ca.* Jettenia in our study are enriched by anammox bioreactors, within which they are often found to reside in granules, which are compact aggregates with a nearly spherical external appearance [79]. Consistent with the niche preference of *Ca.* Jettenia bacteria [80], the genes (*cpaABCEF* and *tadBC*) for pilus assembly and adherence are solely present (Fig. S9). Likewise, multiple acquired genes (*gdh*, *waaD*, *rgpA*, *wbtD*, *mshA*, *epsH*, *mapP*, *hypBA*, *fbaB* and *hddC*) encoding enzymes for carbohydrate utilizations might provide the ability to obtain energy from other microorganisms, which agrees with the potential commensalism between *Ca.* Jettenia and its neighboring heterotrophs [81, 82]. On the other side, the potential loss of chemotaxis genes (*cheABCDR*) (Fig. S9) might be ascribed to the stable and anoxic microenvironment inside granules and hence unnecessity of rapid response to changes in the environmental cue availability.

The genus *Ca.* Brocadia contains the largest genomes and is presumably most physiologically diverse compared to the other genera, which include but are not limited to its acetate oxidization ability, unusual autofluorescent ability [83] and hydroxylamine-dependent anammox process [84]. Among the potentially acquired genes are *ackA*, *ykhA*, *pta* for acyl-CoA assimilation [85], and genes mainly involved in the transportation of urea and heme (*UTP* and *AglD2*) respectively (Fig. S9). The *Ca.* Brocadia also acquired a complete cassette of type III CRISPR-Cas systems (Fig. S9), which could protect them against invading viruses and plasmids [86]. In addition, genes involved in flagellar assembly, in particular, the genes encoding hook (*flgABCDEFGHIJKL*), filament (*fliCDEHKLQW*) and chemotaxis genes (*cheABCDR*) are present in the genomes of *Ca.* Brocadia (Fig. S9).The coexistence of chemotaxis and flagella-encoding genes implies the potential to move in response to environmental stimuli [87]. This is consistent with previous reports that in case nitrogen supply was limited *Ca.* Brocadia bacteria could move in the direction of higher concentrations of nitrogen [88].

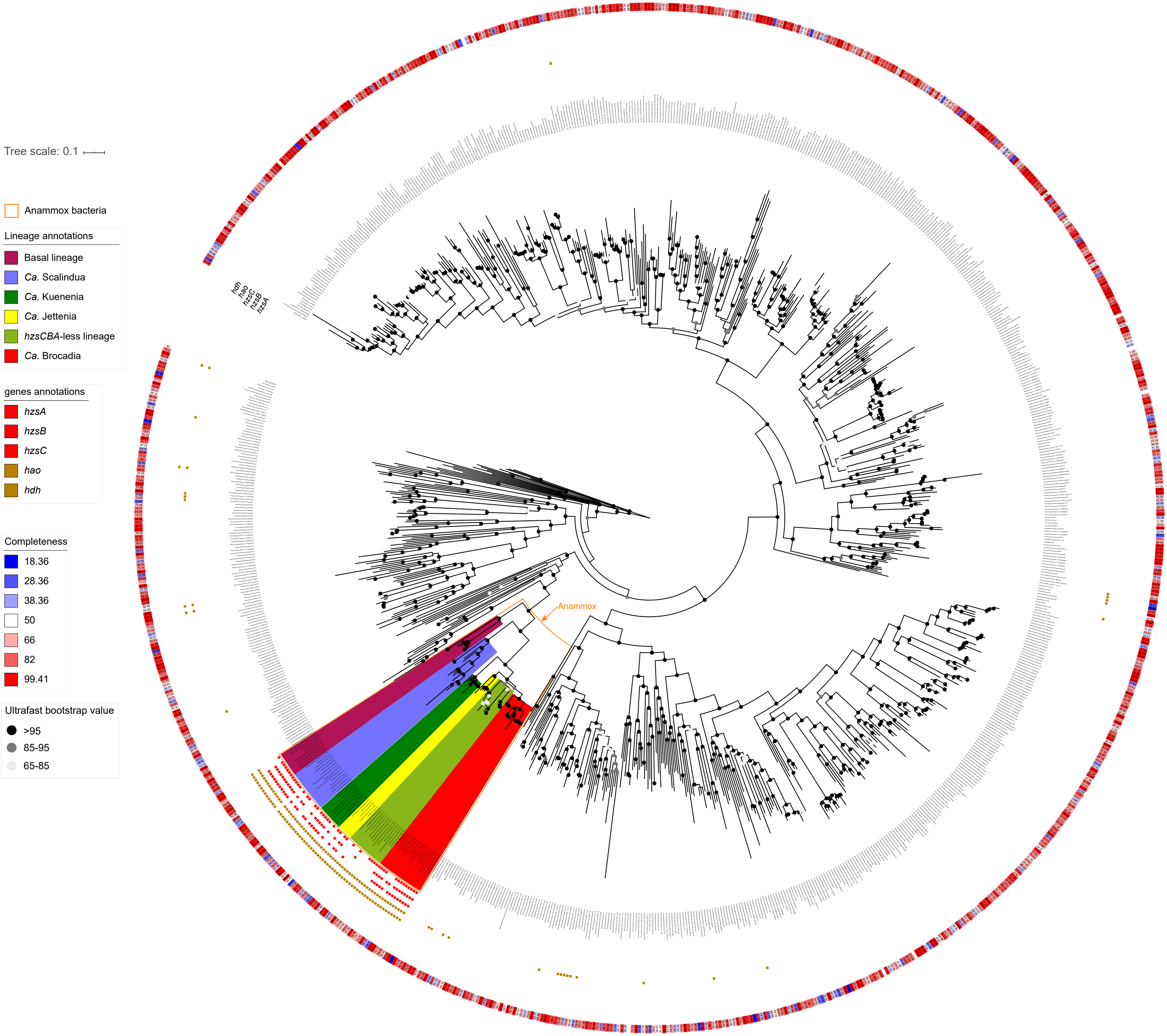


**Figure S1**. The phylogenomic tree of the 881 genomes from the phylum Planctomycetes and six genomes from the phylum Verrucomicrobia which are taken as the outgroup. The tree is built with the 120 bacterial proteins (bac120) widely used for tree inference of prokaryotes. The genome completeness estimated by checkM is visualized with the gradient color strip at the outermost ring. The squares arranged as rings around the tips indicate the presence of key genes of anammox including *hzsCBA* (red), and *hdh* or *hao* (yellow). Genomes without any *hzsCBA* are not annotated.

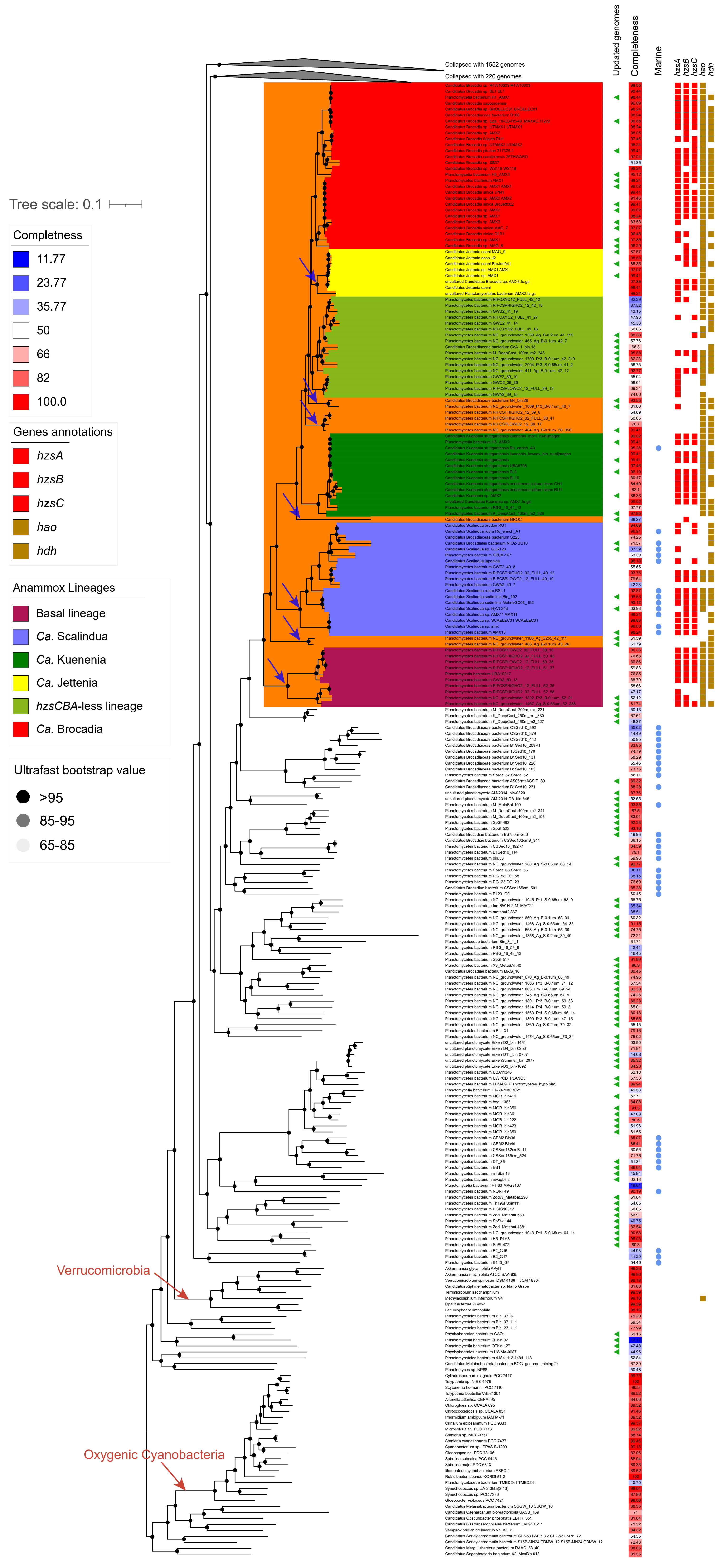


**Figure S2**. The phylogenomic tree built with the same method as Figure S1 using the latest Planctomycetes genomes (Genome set 2; see Dataset S1.1). The green triangle besides the tip label denotes updated genome (those that are not present in Figure S1). The color strip next to it indicates the genome completeness estimated by checkM. The blue filled circles indicate the genomes sampled from marine habitats. The blue arrows represent multiple potential losses of the ability of utilizing NO as the sole electron acceptor if the LCA is assumed to be NO-denpendent anammox bacteria.

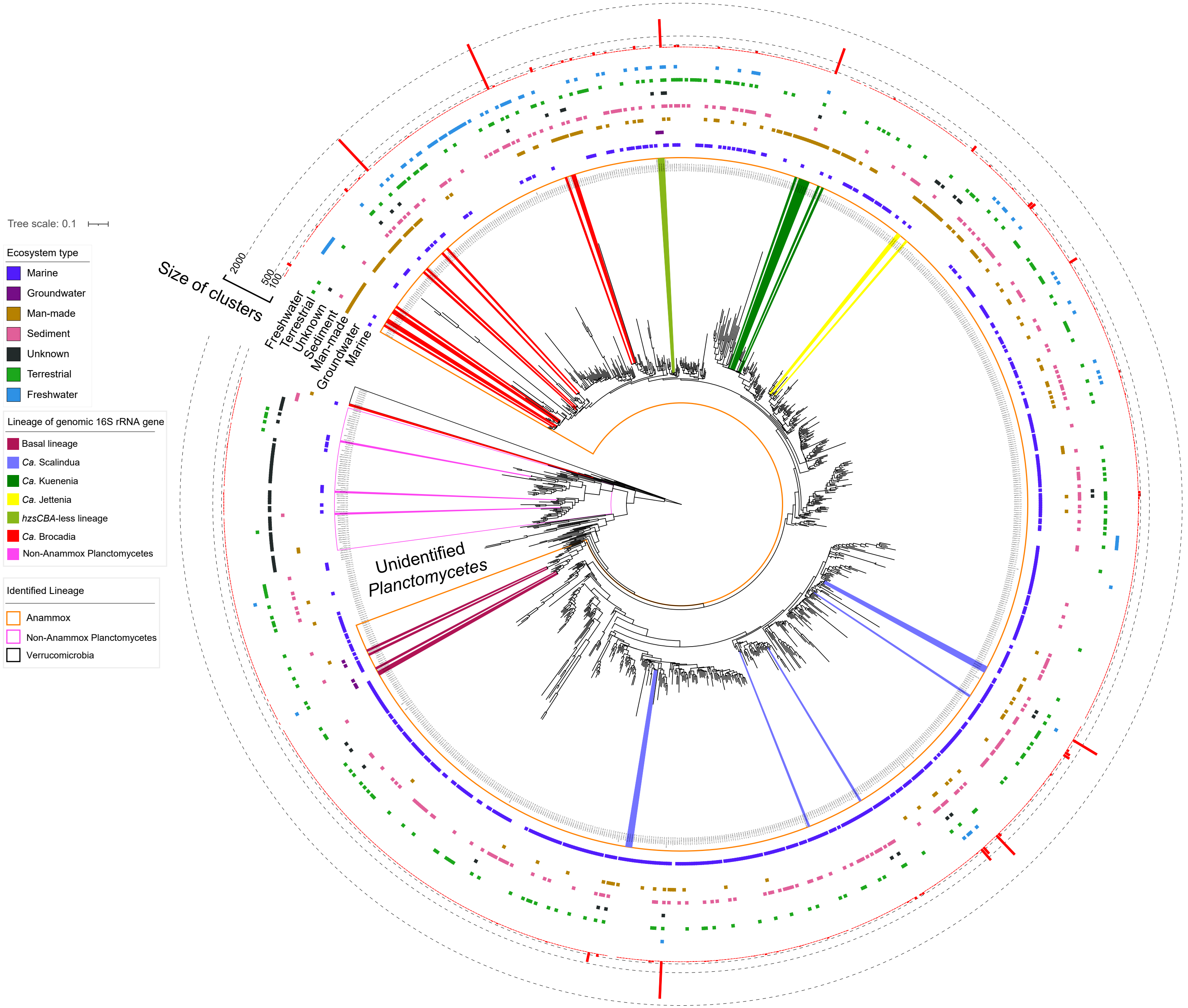


**Figure S3**. The phylogenetic tree built with 16S rRNA genes from the 881 Planctomycetes and the six Verrucomicrobia genomes and representative 16S rRNA sequences affiliated with the class Brocadiae (both anammox and non-anammox bacteria included) from the SILVA database. The tree is rooted with Verrucomicrobia (black box). The tips with colors are 16S rRNA genes annotated from genomic sequences as reference. The height of the red bars represents the size of clusters of 16S rRNA genes. Note that sediments from mangrove, estuary, intertidal zone and salt marsh are classified as marine. The raw isolation sources are provided in Dataset S1.3 and online repository (see Data and Code availability).

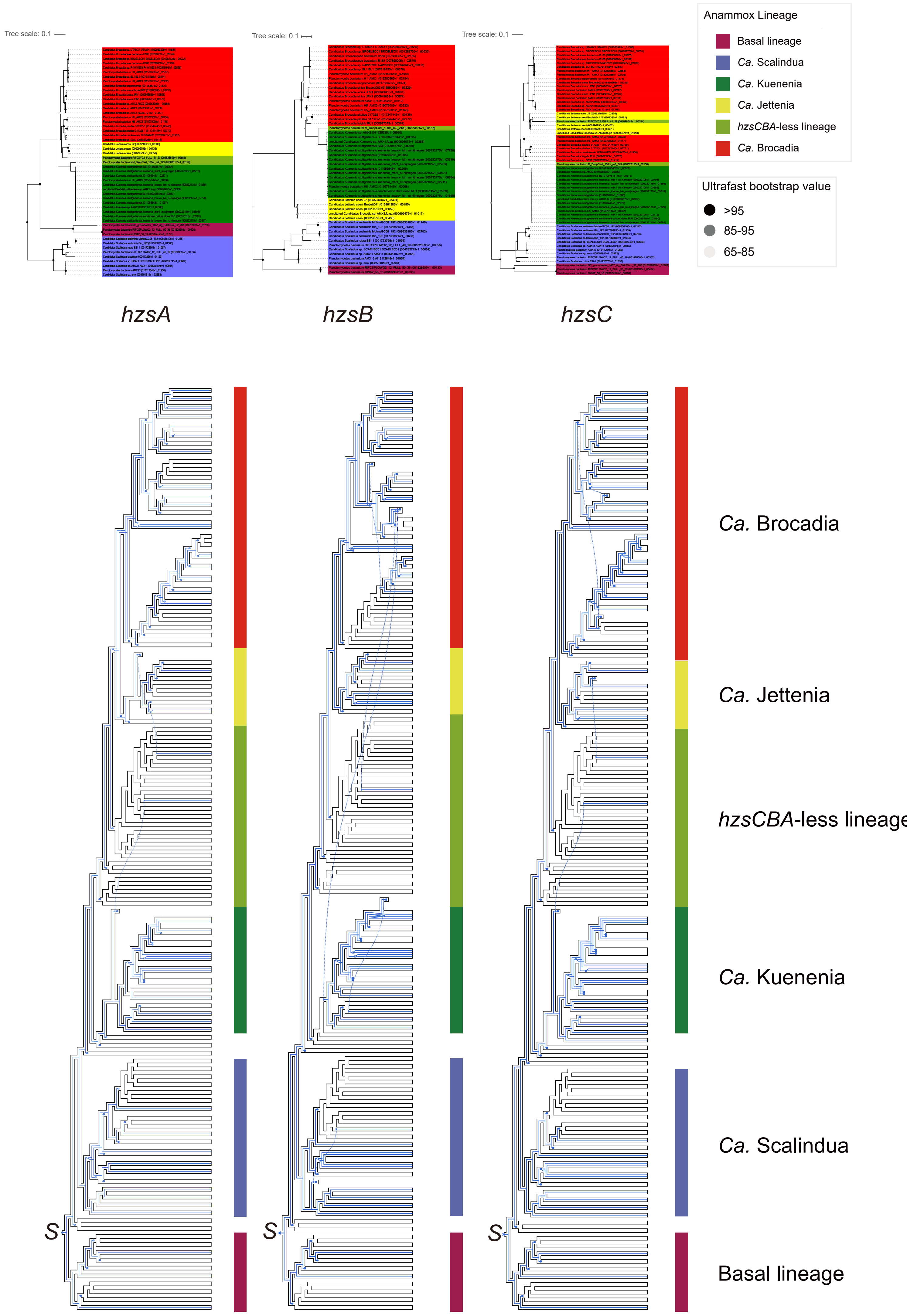


**Figure S4**. The gene phylogenies and reconciliation results of *hzsCBA*. Shown on the top of the figure are the phylogenetic trees of HzsA, HzsB, and HzsC, which are rooted by the minimal ancestor deviation (MAD) method. The phylogenies with inferred evolutionary events at the bottom of the graph are the visualization of gene reconciliation results which are inferred using GeneRax. The blue lines within the branches indicate the inference of the evolutionary history of corresponding genes. The S nodes indicate the origin of corresponding genes. The X symbol indicates an inferred loss. The square symbol indicates an inferred duplication event. The dotted line indicates an inferred gene transfer event.

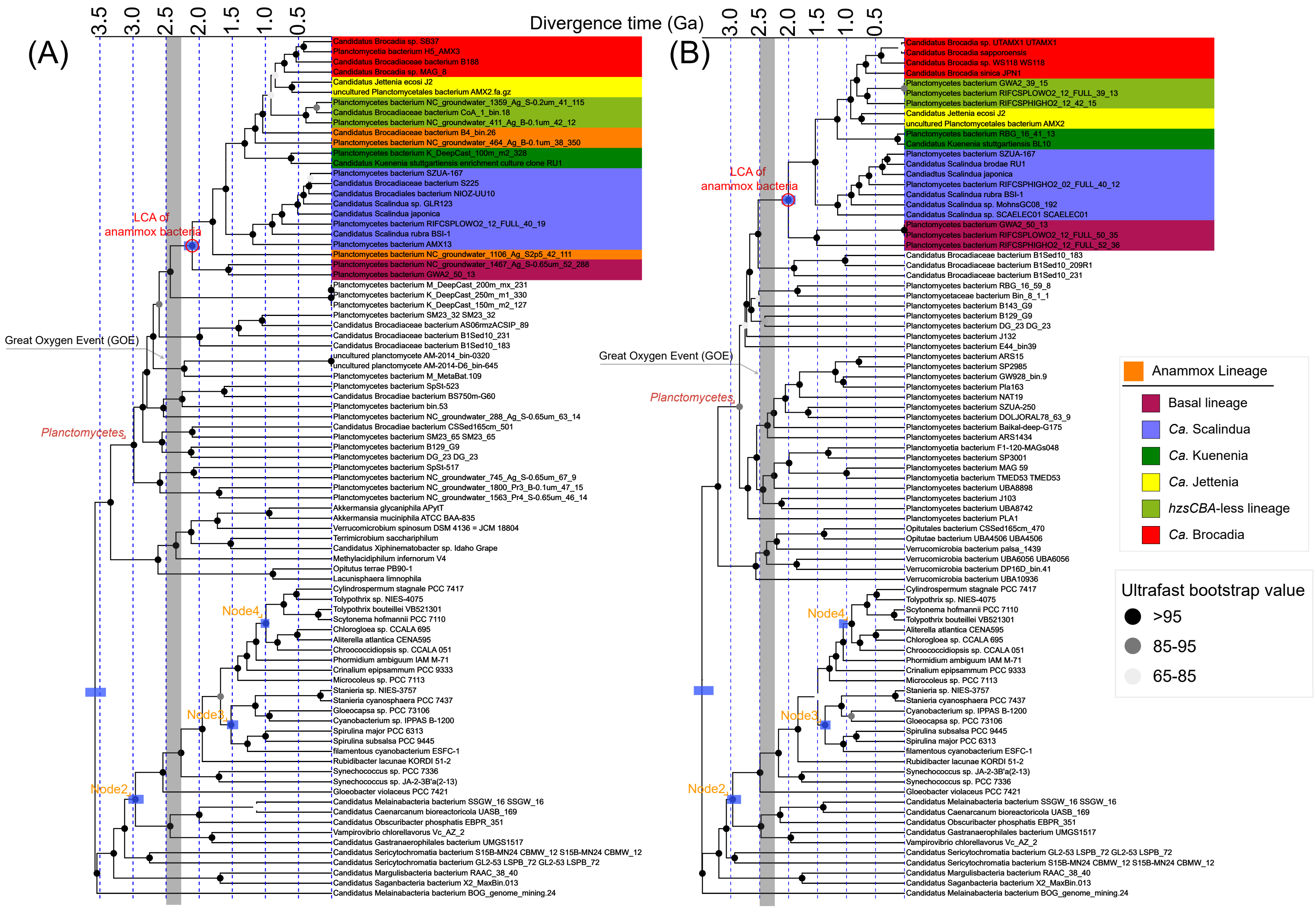


**Figure S5**. The evolutionary timelines of anammox bacteria from two repeat analyses. (A) topology inferred from the analysis based on the Genome set 2 (Dataset S1.1) (B) topology based on Genome set 1 (Datasets S1.1 & S1.2) but with the LG+C60+G model. Two chronograms were individually estimated based on the calibration set C1 and independent rate model (see Supplementary Text section 3). The calibration constraints used within the phylum Cyanobacteria are marked with orange texts: the LCA of Planctomycetes and Cyanobacteria (Root; smaller than 4.5 Gyr), the total group of oxygenic Cyanobacteria (Node 2; larger than 2.32Gyr and smaller than 3.0 Gyr), the total group of Nostocales (Node 3; larger than 1.6Gyr), and the total group of Pleurocapsales (Node 4; larger than 1.7Gyr). The vertical grey bar represents the period of the GOE from 2,500 to 2,320 Ma. The blue bars on the four calibration nodes and the last common ancestor (LCA) of anammox bacteria represent the posterior 95% highest probability density (HPD) interval of the posterior time estimates.

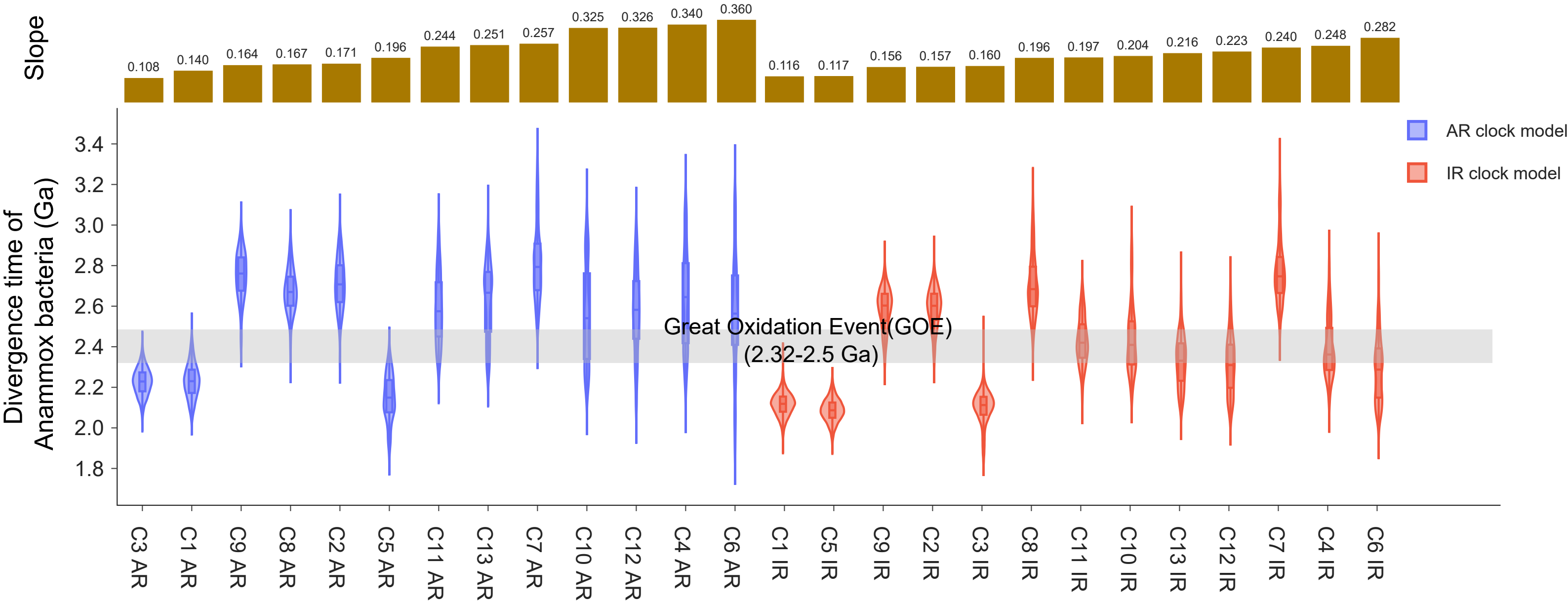


**Figure S6**. The divergence times of anammox bacteria estimated using MCMCTree under different calibration sets (C1-C13). The calibration sets are ordered by the slope inferred from the infinite-site plot (see Supplementary Text section 3.4). The horizontal grey bar represents the great oxygen event (GOE) from 2.5 to 2.32 Ga. The detailed constraints of calibrations and time estimates are provided in Dataset S2.1.

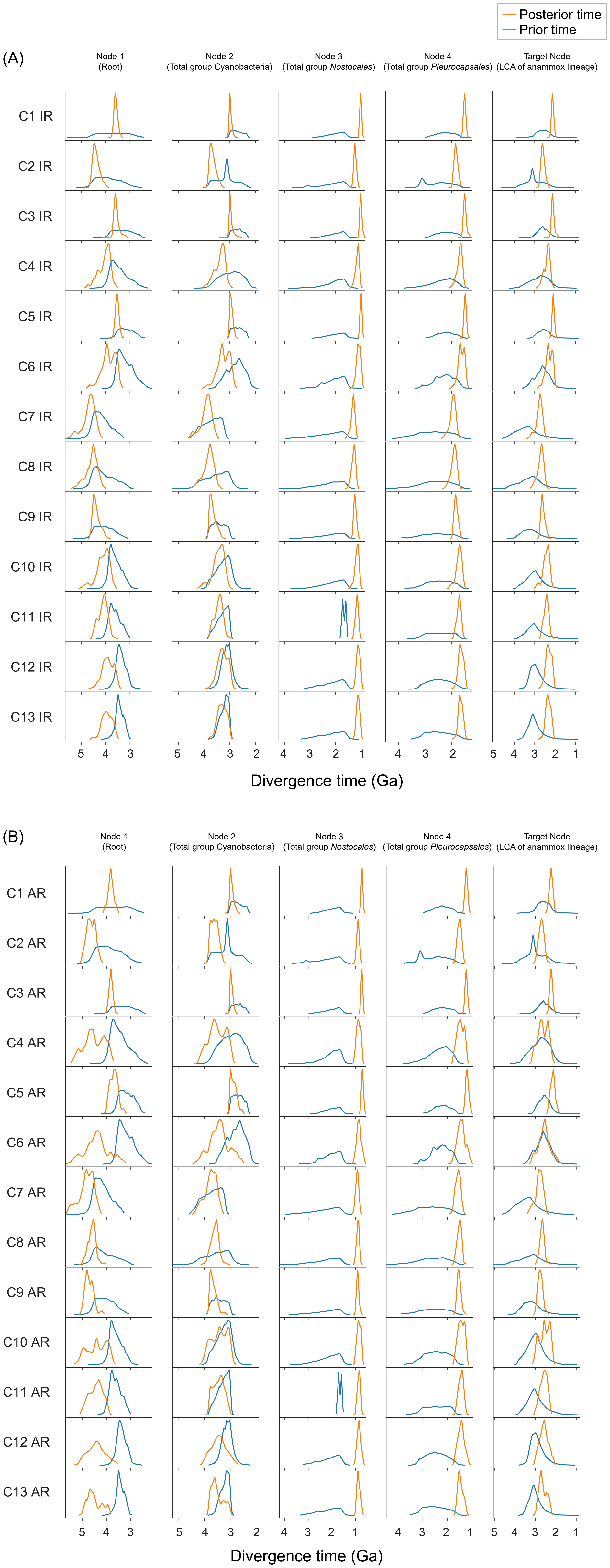


**Figure S7**. The densities of posterior (orange line) and prior (blue line) divergence times of the four calibrated nodes and the target node (LCA of anammox lineage) with 26 different sets of time constraints. IR (A) and AR (B) in the dating scheme ID (on the x-axis; Dataset S2.1) represent auto-correlated rates model and independent rates model, respectively.

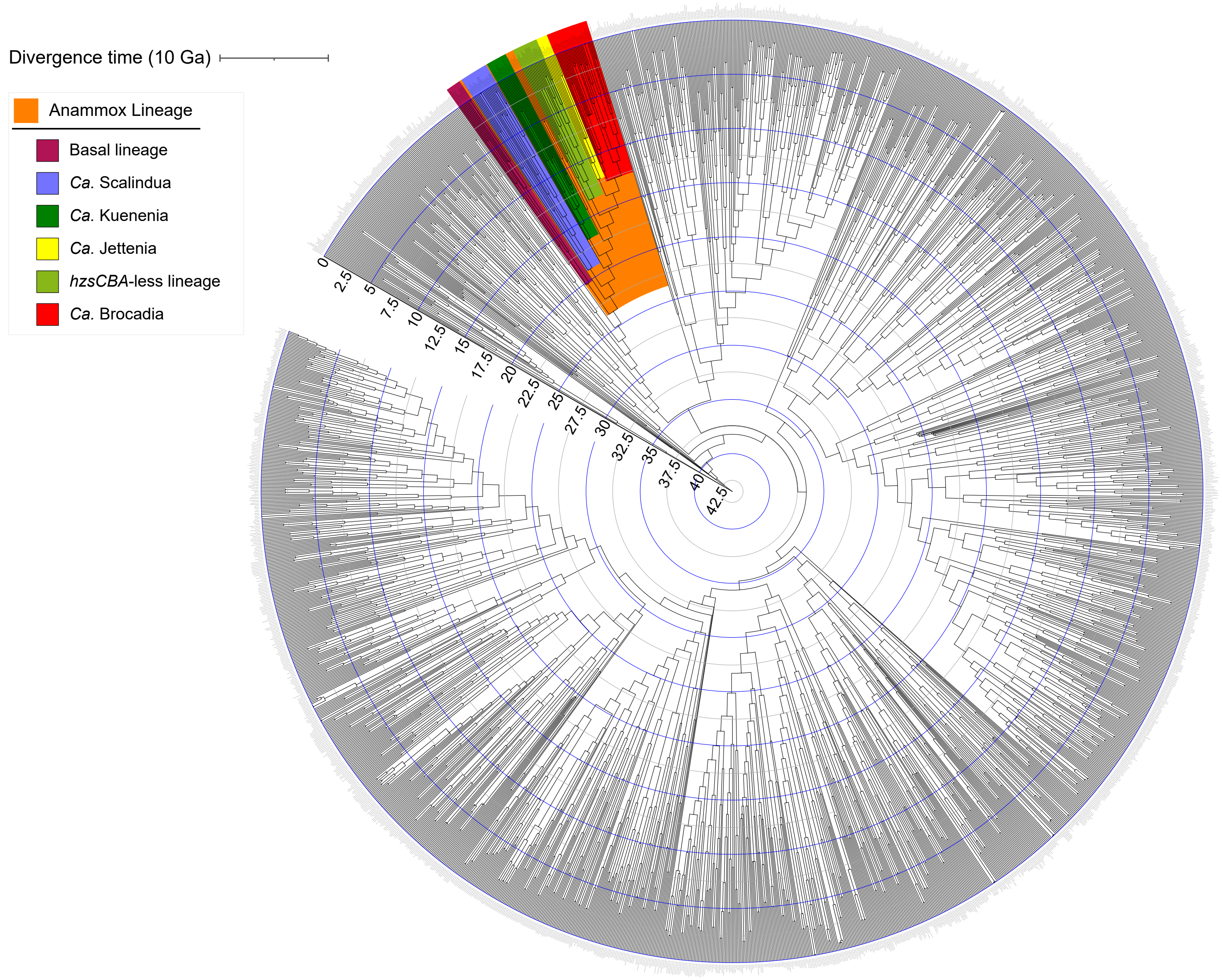


**Figure S8**. The radial chronogram was generated based on the phylogenomic tree shown in Figure S2 using likelihood-based dating analysis with IR model.

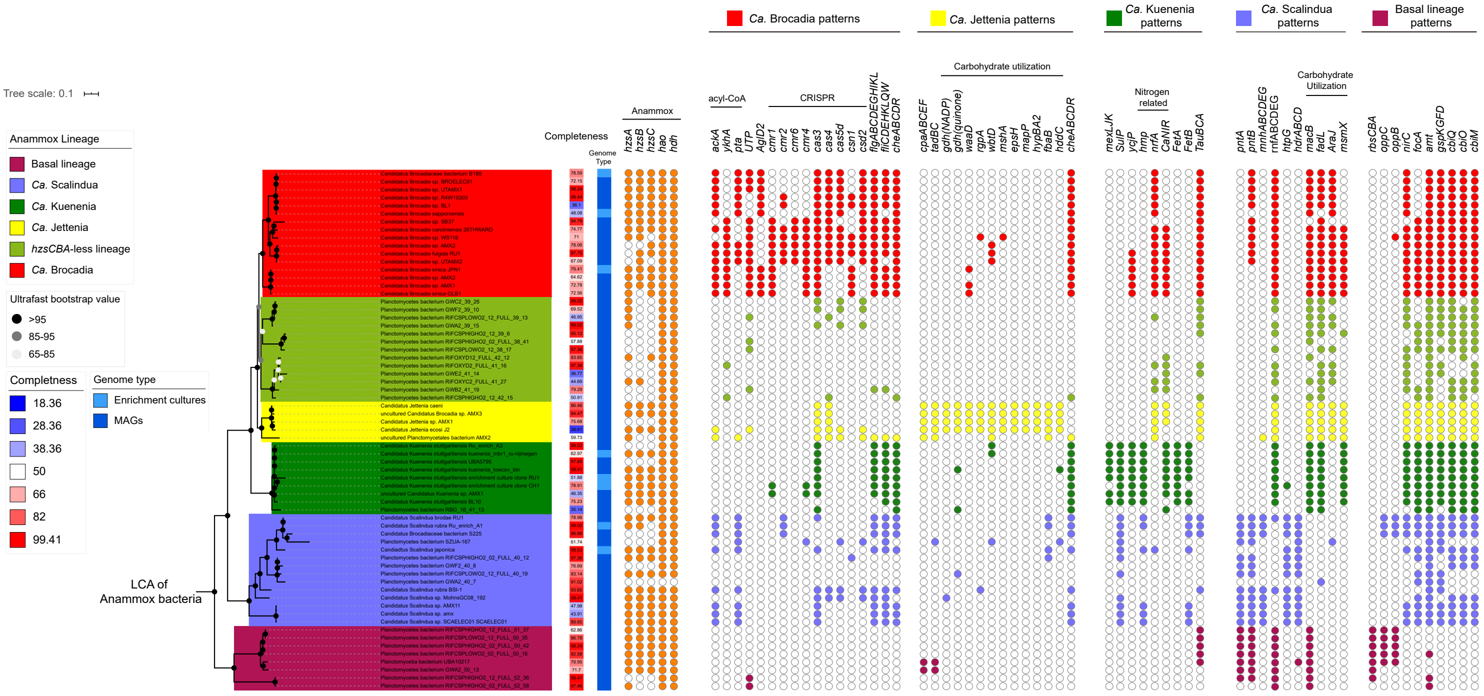


**Figure S9**. The phyletic pattern of genes that show statistical differences in the comparison between each anammox genus and other anammox bacteria using multiple fisher's exact tests. The *hzsCBA*-less lineage is not shown because few differences in the gene phyletic pattern between this and other anammox bacteria lineages are observed. The phylogenomic tree on the left was pruned to show the anammox lineage only from Figure S1. The estimated genomic completeness using checkM was visualized and labelled with gradient color strip. The other color strip represents the type of genomic sequences including metagenome-assembled genomes (MAGs) and whole-genome sequencing (WGS) of enriched culture samples (a few anammox bacteria) used in our study. The filled and empty circles represent the presence and absence of particular genes in corresponding genomes, respectively. *ackA*, acetate CoA; *ykhA*, acetate CoA acyl-CoA thioester hydrolase; *pta*, acetate CoA; *UTP*, urea transporter; *AglD2*, putative heme transporter; *cmr1, cmr2, cmr6, cmr4, cas3, cas4, cas5d, csn1, csd2*, CRISPR-associated protein; *flgABCDEGHIKL*, flagella basal, hook and assembly protein; *fliCDEHKLQW*, hook and assembly protein; *cpaABCEF*, pilus assembly; *tadBC*, adherence; *gdh*, (NADP) glucose 1-dehydrogenase; *gdh*, (quinone) glucose dehydrogenase; *waaD*, UDP-glucose alpha-1,2-glucosyltransferase; *rgpA*, rhamnosyltransferase; *wbtD*, galacturonosyltransferase; *mshA*, D-inositol-3-phosphate glycosyltransferase; *epsH*, glycosyltransferase; *mapP*, maltose 6'-phosphate phosphatase; *hypBA2*, beta-L-arabinobiosidase; *fbaB*, fructose-bisphosphate aldolase; *hddC*, mannose-1-phosphate guanylyltransferase; *cheABCDR*, chemotaxis protein; *mexLJK*, multidrug efflux system; *SulP*, Sulfate Permease; *ycjP*, multiple sugar transport system permease; *hmp*, nitric oxide dioxygenase; *nrfA*, ammonia-forming nitrite reductase subunit A; *CaNIR*, calcium-dependent multiheme nitrite reductase; *FetAB*, ABC transport for iron; *TauBCA*, nitrate/nitrite transporter; *pntAB*, $H^+$-translocating NAD(P) transhydrogenase subunits alpha and beta; *mnhABCDEG*, multicomponent $Na^+/H^+$ antiporter; *htpG*, heat shock protein; *hdrABCD*, heterodisulfide reductase; *macB*, macrolide transport; *fadL*, long-chain fatty acid transporter; *AraJ*, multidrug resistance protein; *msmX*, multiple sugar transport system ATP-binding protein; *rbsCBA*, ribose transporter; *oppBC*, oligopeptide transport system; *nirC*, nitrite transporter; *focA*, formate transporter; *amt*, ammonium transporter; *gspKGFD*, general secretion pathway protein; *cbiQ, cbiO, cbiM,* cobalt/nickle transport system protein.
